# *In silico* and *in vitro* studies suggest epigallocatechin gallate (EGCG), a polyphenol in green tea, can bind and modulate the aggregation and cytotoxicity of the full-length TDP-43 protein implicated in TDP-43 proteinopathies

**DOI:** 10.1101/2023.12.22.573011

**Authors:** Vini D. Meshram, Ramkumar Balaji, Preethi Saravanan, Yashashwini Subbamanda, Waghela Deeksha, Akarsh Bajpai, Himanshu Joshi, Anamika Bhargava, Basant K. Patel

## Abstract

Misfolding and aggregation of TDP-43 protein are implicated in several proteinopathies like ALS and FTLD. Extracellular TDP-43 is also proposed to propagate in a prion-like pathogenic manner to the neighbouring cells. Here, using turbidity and sedimentation assay, we show that a polyphenol in green tea, epigallocatechin gallate (EGCG), can inhibit the *in vitro* aggregation of the full-length TDP-43 protein. Furthermore, Alexa-Fluor-labelled TDP-43 protein failed to show aggregates in the presence of EGCG in fluorescence microscopy. Also, AFM imaging revealed that EGCG co-incubation with TDP-43 allows formation of only small oligomers in contrast to the larger TDP-43 aggregates formed otherwise. A physical binding of EGCG with TDP-43 was observed using triphenyl tetrazolium chloride (TTC) staining and isothermal titration calorimetry (ITC). ITC also revealed a high-affinity binding site for EGCG on TDP-43 with a K_d_ value of 7.8 µM and a binding free energy of -6.9 kcal/mol. Furthermore, *in silico* molecular docking and molecular dynamic simulation (MDS) studies using different available structures of the N-terminal, RRM1-2 and C-terminal domains of TDP-43, predicted a preferable and stable binding of EGCG to the structure of the aggregation prone C-terminal domain (CTD) (PDB ID:7KWZ). Also, EGCG complexed with CTD of TDP-43 yielded a negative ΔG value of -20.29 kcal/mol using MM-PBSA analysis of the MDS data thereby further suggesting a stable complex formation. Also, in MDS, EGCG interacted with the amino acids Phe-313 and Ala-341 of TDP-43, which were previously projected to be important for the recruitment of monomers for the amyloid formation by CTD, thereby suggesting a possible mechanism of EGCG’s inhibition of the TDP-43 aggregation. Notably, while the *in vitro*-made aggregates of full-length TDP-43 caused mild cytotoxicity to the HEK293 cells, the small oligomers of TDP-43 formed in presence of EGCG did not. In totality, EGCG can *in vitro* interact with TDP-43 and inhibit its aggregation, possibly *via* interaction with the amyloidogenic domain, thereby preventing it from assuming cytotoxic conformations. As EGCG is a natural molecule, it could be relevant to the therapeutic quest against the TDP-43 proteinopathies.

## 1. Introduction

The misfolding and proteinopathy of TDP-43 is associated with several diseases such as amyotrophic lateral sclerosis (ALS) and frontotemporal lobar degeneration (FTLD) [1]. In TDP-43 proteinopathies, TDP-43 which is normally a nuclear protein undergoes mis-localization to the cytoplasm where it forms aggregates [1]. The TDP-43 protein comprises of a well folded N-terminal domain and two RNA recognition motifs (RRM1 and RRM2) which are very well conserved, and an aggregation-prone intrinsically disordered C-terminal domain (CTD) harbouring a prion-like glutamine/asparagine-rich sequence [2]. TDP-43’s CTD promotes its liquid-liquid phase separation (LLPS) which can eventually lead to solid-like phase transition [1,3]. Several studies using the TDP-43 aggregation models such as the *Drosophila* model system show a correlation between the aggregation of TDP-43 and cytotoxicity [4]. Notably, prion-like propagation *via* cell-to-cell transmission leading to disease progression has been reported for the α-synuclein protein [5]. In fact, the progression of ALS is characterised by a regional spreading of the symptoms [6] and a prion-like propagation of TDP-43 *via* intercellular spread has also been hypothesised for the disease spreading. Some findings support this hypothesis that the pathogenic aggregates of TDP-43 have prion-like properties including cell-cell transfer and the ability to seed the intracellular TDP-43’s aggregation [7,8].

Recently, the full-length TDP-43 aggregates were deciphered to have an amyloid-like core, inherently masked by the flanking structured domains, but could be revealed upon partial proteinase K proteolysis of the full-length TDP-43 aggregates [9]. In view of the pathological relevance of the TDP-43 aggregates, identification of modulators of its aggregation may be a potential therapeutic strategy against TDP-43 proteinopathies. Such studies have been carried out *in vitro* using different model systems and targeting the aggregation of different domains of TDP-43 using ligands such as nTRD22, poly-ADP-Ribose (PAR), the drug mitoxantrone and 8-hydroxyquinoline-based small molecules [10–13]. Additionally, *in silico* approaches towards design of inhibitors towards TDP-43 aggregation are also being strategized [14,15]. Previously, we found an imidazole containing acridine derivative, AIM4, that could prevent the *in vitro* aggregation of a C-terminal fragment of TDP-43, termed TDP-43^2C^ [16]. AIM4 possess a planar fused tricyclic ring and was shown to interact favourably to the CTD leading to inhibition of the LLPS and the aggregation of TDP-43^2C^ [16,17].

The organic molecule, epigallocatechin Gallate (EGCG) is a bioactive component found in the commonly used beverage, green tea. It belongs to the class of polyphenols with a large number of active phenolic hydroxyl groups linked to four core rings **(Figure 1)** [18]. Previous studies have reported that EGCG has anti-aggregation properties and can modulate the misfolding of certain disease-causing proteins and prions such as tau, α-synuclein and amyloid-β protein [19]. Studies on mutant-SOD1 expressing mice have suggested a disease modifying role of EGCG against ALS [20].

**Figure 1:**
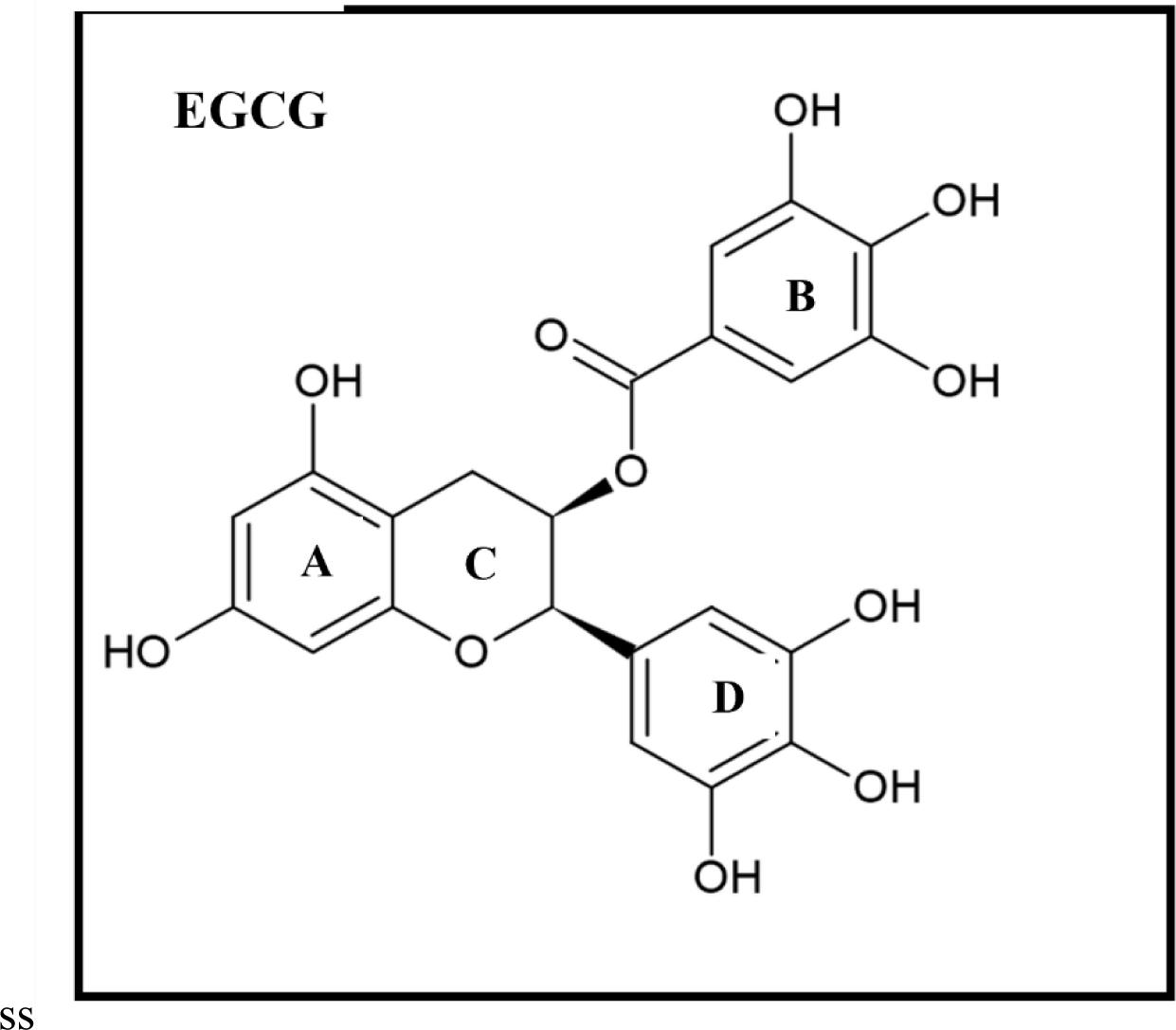
Structure of green tea polyphenol Epigallocatechin Gallate (EGCG).

In another study upon EGCG treatment to the 293T cells that were expressing TDP-43, it was found that a TDP-43’s C-terminus mediated aggregation of the natively folded TDP-43 occurs into non-cytotoxic functional oligomers thereby inhibiting its pathological-like degradation [21]. However, a direct physical binding of EGCG with the TDP-43 protein has not been investigated yet. Therefore, in this study, we have investigated the inhibitory potential of EGCG against the *in vitro* aggregation of the full-length TDP-43 protein using tools such as turbidity assay, fluorescence microscopy and atomic force microscopy. Furthermore, we examined the physical interaction of EGCG with TDP-43 using TTC staining and isothermal titration calorimetry. Further, cell viability assays were performed to examine the relative cytotoxicities of the TDP-43 aggregates formed in the presence or absence of EGCG treatment. Finally, to explore the probable binding site(s) and residues involved in the interaction of EGCG with TDP-43, *in silico* molecular docking and molecular dynamics simulation studies were performed using available structures of the various TDP-43 domains.

## 2. Materials and methods

### 2.1. Plasmids and strains

For recombinant protein expression, *Rosetta 2 (DE3) E. coli* strain and the plasmid *pET15b-His-TDP-43* encoding the full-length TDP-43 protein (a kind gift of Prof. Yoshiaki Furukawa, Keio University), were used [22].

### 2.2. Expression and purification of recombinant TDP-43

Recombinant expression and purification of N-terminal His-tagged TDP-43 was carried out as done previously [16,17,23,24]. Briefly, *Rosetta 2 (DE3) E. coli* competent cells (Novagen, USA) pre-transformed with the *pET15b-His-TDP-43* plasmid were induced for 4 h with 1mM isopropyl-β-D-1-thiogalactopyranoside (IPTG) for protein expression and cells were harvested and lysed by ultra-sonication in lysis buffer (6 M guanidine HCl (GuHCl) in phosphate buffered saline (PBS) pH 7.5 and 1 mM of PMSF). The pre-cleared lysate was loaded onto a Ni-NTA agarose affinity chromatography column pre-equilibrated with 6 M GuHCl in PBS pH 7.5. Non-specifically bound proteins were washed out with buffer (6 M GuHCl in PBS pH 7.5 and 10 mM imidazole). Subsequently, bound TDP-43 protein was eluted with elution buffer (250 mM imidazole and 6 M GuHCl) in fractions of 500 µL. Protein homogeneity was determined on 10% SDS-PAGE.

### 2.3. Assays for in vitro aggregation of TDP-43 and inhibition by EGCG

#### 2.3.1. Turbidimetry

As aggregation of protein causes an increase in solution turbidity, the effect of EGCG on TDP-43 turbidity was examined by recording absorbance at 450 nm [25,26]. For this, TDP-43 purified in 6 M GuHCl was first diluted 10-fold in PBS pH 7.5 resulting in the precipitation of the protein. The precipitate was pelleted by centrifugation at 14,000 rpm for 10 minutes at 4°C, the supernatant was discarded to remove GuHCl and the protein stock solution was prepared by re-suspending the pellet in 6 M urea in 10 mM Tris pH 7.5. The protein stock was diluted 20-fold with 10 mM Tris, pH 7.5 to allow for rapid refolding of the protein and then centrifuged at 20,000g for 30 minutes at 4°C. The concentration of refolded protein in supernatant was determined by measuring the absorbance at 280 nm. The refolded protein diluted to 10 µM in aggregation buffer (50 mM Tris, 50 mM NaCl, and 1 mM DTT) was incubated in the presence or absence of EGCG (protein: EGCG ratio of 1:2) at 37°C with shaking and absorbance at 450 nm was recorded at different incubation times up to 6 h.

#### 2.3.2. Sedimentation Assay

The effect of EGCG on the particulate nature of TDP-43 was analysed by examining the relative difference in the amount of insoluble aggregated protein in presence or absence of EGCG upon sedimentation. For this, refolded TDP-43 was diluted to 10 µM in aggregation buffer (as mentioned in section 2.3.1) and incubated in the presence and absence of 20 µM EGCG at 37°C with shaking. The proteins samples were fractionated using previously used methods [27]. Unfractionated sample (total, T), the supernatant (S) and the resuspended pellet (P) from the fractionated samples were treated with Laemmli reducing buffer and heated at 99°C for 10 minutes. Then, all fractions were spotted on to a PVDF membrane, followed by staining with Coomassie staining solution and destaining to visualize the protein content.

#### 2.3.3. Analysis of aggregate formation by Alexa Fluor-labelled TDP-43 by fluorescence microscopy

To visualize the *in vitro* formed aggregates of TDP-43 in the presence and absence of EGCG, TDP-43 was labelled with free cysteine reacting fluorescent dye Alexa Fluor Maleimide-488 as done in previous studies [16,24]. For this, the protein mixture of unlabelled TDP-43 and Alexa Fluor-labelled TDP-43 maintained at the ratio of 1:10, was diluted to a final protein concentration of 100 µM in aggregation buffer (PBS pH 7.5, 2 M urea and 500 µM DTT) in the absence and presence of EGCG (protein: EGCG ratio of 1:2) at 37 °C with shaking, and observed for any fluorescent aggregate structures at 1 h, 2 h, 4 h and 12 h under Leica DM2500 fluorescence microscope using the GFP filter. Images were processed using the ImageJ software [28].

### 2.4. Morphological analysis of TDP-43 aggregates by atomic force microscopy

Size and morphology of *in vitro* formed TDP-43 aggregates in the presence or absence of 200 µM EGCG were analysed by atomic force microscopy. 10 µL of each sample was placed on a freshly cleaved mica sheet and analysed by multimodal scanning probe microscope (Park Systems) in tapping mode [23]. The images were analysed using XEI software (Park Systems).

### 2.5. TTC-Glycine Assay for EGCG detection

EGCG reduces tetrazolium salts such as 3-(4, 5-dimethylthiazol-2-yl)-2, 5-diphenyltetrazolium bromide (MTT) or nitro blue tetrazolium (NBT) and forms a coloured formazan product [29,30]. Here, the tetrazolium salt, triphenyl tetrazolium chloride (TTC), was used as a stain for examining the direct interaction of EGCG with TDP-43. 100 µL of 100 µM TDP-43 pre-incubated with 100 µM EGCG were subjected to first round of dialysis (15 h) against 500 mL of PBS pH 7.5 followed by a second round of 24 h dialysis against fresh 500mL of PBS pH 7.5.at 4°C using a dialysis membrane with a MWCO of 3.5 kDa. Post-dialysis, EGCG-bound protein along with the control samples such as dialysed protein without EGCG, dialysed EGCG and un-dialysed EGCG (100 µM) were incubated under dark conditions with 0.24 mM TTC in 2 M KOH-Glycine, pH 10.0 in microplate wells for 2 h. Following this, the TTC-reduced red-formazan product was quantified spectrophotometrically by recording absorbance at 500 nm. Simultaneously, 10 µL of the dialysed samples were also spotted onto PVDF membrane followed by addition of 0.24 mM TTC in 2 M KOH-Glycine. The intensity of red-formazan product developed due to the presence of EGCG was then quantified by densitometry using the ImageJ software [28].

### 2.7. MTT cell viability assay

HEK293 cells were seeded in 24 well plates (approximately 40000 cells / well) and incubated in CO_2_ incubator for 24 h. Aggregates of the recombinantly purified full-length TDP-43 were prepared in absence and presence of 200 µM EGCG by incubating 100 µM TDP-43 in the aggregation buffer (2 M urea in PBS pH 7.5) at 37°C with shaking for 1 h. Post this, cells were treated with samples containing either the aggregation buffer or 0.5 µM TDP-43 aggregates or buffer + EGCG or 0.5 µM TDP-43 pre-incubated with EGCG. Subsequently, 72 h post treatment, fresh media was replaced and MTT was added and incubated for 4 h. Media was aspirated and 400µL DMSO was added to dissolve the formazan crystals formed by the cells. Absorbance was taken at 540 nm using a plate reader. All data were analysed using GraphPad Prism version 5.0. Unpaired t-test was used between two groups with parametric distribution. For non-parametric data, Mann-Whitney test was used. *p* ≤ 0.05 was considered significant. * denotes *p* ≤ 0.05; ** denotes *p* ≤ 0.001 and *** denotes *p* ≤ 0.0001.

### 2.8. Isothermal Titration Calorimetry

Isothermal titration calorimetry (ITC) experiment for investigating the binding of EGCG to the full length TDP-43 was carried out in LV Affinity ITC (TA Instruments) at 25°C. For ITC studies, TDP-43 purified in the elution buffer was first refolded as mentioned in section 2.3.1. Then, 20 µM of full-length TDP-43 in refolding buffer (0.3 M Urea in 10 mM Tris, pH 7.5) was titrated with 2 mM EGCG also present in the refolding buffer, pH 7.5. Subsequently, 350 µL of TDP-43 was loaded in the sample cell and 200 µL of EGCG was filled in the injection syringe. EGCG was injected into the sample cells with an initial injection of 0.5 µL followed by 20 consecutive injections of 2.5 µL each at 180 s time intervals while stirring at 125 rpm. A background titration was also performed by injecting EGCG into buffer without protein to measure heat changes resulting from dilution of the titrant. Dilution heat of EGCG was subtracted from experimental data and fitted to sequential three-site binding model using NanoAnalyse software and ΔH, ΔS and the binding constant K_d_ were obtained.

### 2.9. In silico molecular docking studies

For *in silico* docking studies, the crystal, NMR, and cryo-EM structures respectively of the N-terminal (RCSB PDB ID: 5MDI) [31]. RRM-1,2 containing domain (RCSB PDB ID: 4BS2) [32] and the C-terminal of TDP-43 (RCSB PDB ID: 7KWZ) [33], were used. The 3D ligand structure of EGCG (PubChem ID: 65064) was obtained from the PubChem library (https://pubchem.ncbi.nlm.nih.gov/). The EGCG structure was docked to various domain structures and amyloidogenic peptide fragments of TDP-43 from the low complexity domain using Autodock-4.2. The docked poses and the non-covalent interactions were visualized for the top docking conformation of the ligand using AutoDockTools (ADT). The inhibitory constants, as predicted by the Autodock-4.2, for the ligands with the top docking scores, were plotted for all the analysed domain structures of the TDP-43 protein with EGCG.

### 2.10. Molecular Dynamic Simulation studies

All-atom MD simulations were performed using GROningen MAchine for Chemical Simulations (GROMACS) v. 21.4 and v.19, a free software listed under the GNU General Public license software [34]. The domain structures of TDP-43 namely 5MDI (N-terminal, aa: 2-80), 4BS2 (RRM1-2, aa: 96-269) and 7KWZ (amyloid fibrils of the C-terminal, aa: 276-414) were simulated with and without EGCG to visualize its binding and complex formation of the individual domains. The all-atom structure obtained from docking was used as the starting conformation, whereas the topology files of the protein were prepared using the CHARMM36 – all atom forcefield and pdb2gmx script. The ligand topology file was created using CHARMM General Force Field (CGenFF) server [35,36]. Subsequently, the combined topology of the complex was created using GROMACS input generator. The complex was solvated in a dodecahedron box water using a pre-defined generic equilibrated 3-point solvent model, with the box edges kept 1.5 nm away from the protein. The solvated system was neutralized by adding Na^+^ or Cl^-^ counter-ions based on the protein using the genion script of GROMACS. Thus, assembled all-atom models contained about 148,000 atoms for complex of 7KWZ with EGCG in water and shown in figure S3. The system containing RRM1-2 domain and N-terminal domain with EGCG in water contained 36,638 atoms and 13,182 atoms respectively **(Supplementary Figure S3)**. The all-atom models were subjected to energy minimization using the steepest descent algorithm to avoid any bad contacts in the build configuration. The energy minimized system was then equilibrated at constant number of atoms, constant pressure (P=1atm) and constant temperature (T=300 K), NPT ensemble, for 1 ns followed by constant number of atoms (N), constant pressure (V) and constant temperature, NVT ensemble for 1 ns. During the NPT and NVT equilibration, harmonic position restraints were applied to the protein and the ligand atoms. Finally, systems were simulated for the production run using an NPT ensemble for either 800 ns or 300 ns by removing the position restraints on the protein and the ligand atoms. A 2-fs integration time step and periodic boundary conditions were applied, and particle mesh Ewald (PME) method over a 12Å resolution grid is used to calculate the long-range electrostatic interaction [37]. 300K temperature and 1 atm pressure were maintained in the simulation using V-rescale [38] and Berendsen [38] coupling method respectively. The trajectories from the production runs were used for further analysis and a clustering of the simulation trajectory is performed using the in-build “gmx cluster” script from gromacs. The non-covalent interactions between the protein and the ligand are obtained using PyContact [39]. The free energy of binding for the ligand and the protein was calculated using Molecular Mechanics Poisson-Boltzmann Surface Area (MM-PBSA) method with gmx_MMPBSA [40] tool that uses the MMPBSA.py script [41]. Visual Molecular Dynamics (VMD) [42], BIOVIA Discovery Studio Visualizer 2021 [43] and PyMOL [44] visualization tools were used for visualizing the interactions in the representative structures from the cluster analysis.

## 3. Results

### 3.1. Epigallocatechin Gallate (EGCG) inhibits the in vitro aggregation of TDP-43

Aggregation of the full-length TDP-43 protein and the effect of EGCG on the aggregation was studied by recording the changes in the protein solution turbidity [25,26]. Prior to the aggregation assays, the purified TDP-43 present in denaturing buffer was buffer exchanged to a refolding buffer containing 0.3 M urea, 10 mM Tris HCl pH 7.5 and this refolded TDP-43, when characterized using CD spectroscopy, manifested secondary structural content similar to as reported previously by the other groups for the refolded full-length TDP-43 protein [45] **(Supplementary Figure S1)**. For the turbidity assay, 10 µM refolded protein was incubated in the absence or presence of 20 µM EGCG in the aggregation buffer and changes in turbidity of the solutions was recorded with time by measuring absorbance at 450 nm. Notably, the TDP-43 samples incubated in the absence of EGCG showed an increase in the solution turbidity with time **(Figure 2a)**. On the contrary, TDP-43 samples co-incubated with EGCG at stoichiometric ratio of 1:2 did not manifest any increase in the solution turbidity over time **(Figure 2a)**. This suggests of inhibition of the aggregation of TDP-43 by EGCG. Furthermore, when the effect of EGCG on the aggregation of TDP-43 was also studied under mild denaturing conditions by incubating 100 µM TDP-43 with 200 µM of EGCG in aggregation buffer containing 2.5 M urea, EGCG showed similar inhibitory effect on the aggregation of TDP-43 **(Supplementary Figure S2a)**.

**Figure 2:**
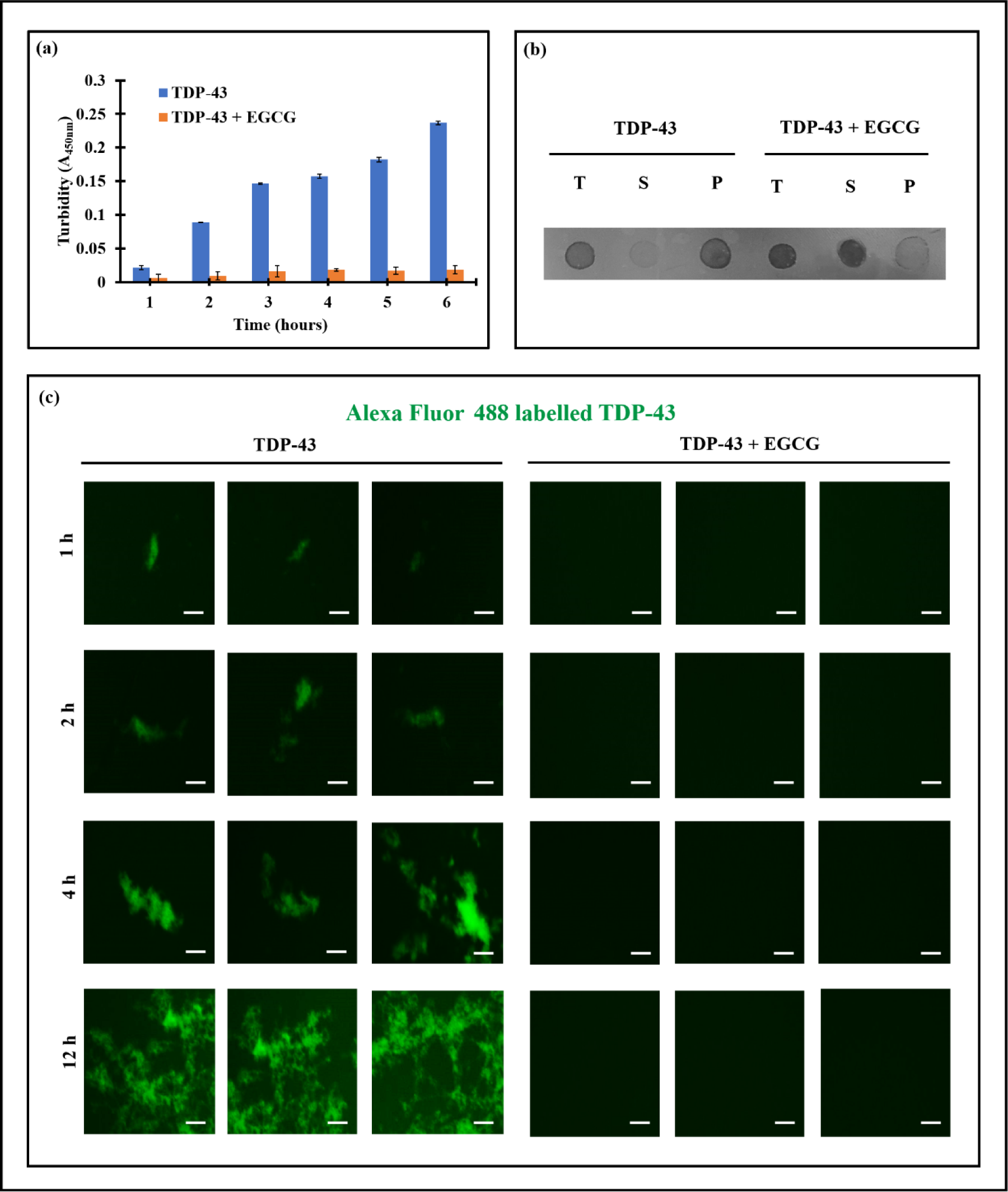
EGCG can inhibit the *in vitro* aggregation of full-length TDP-43. (a) Changes in the turbidity of the refolded TDP-43 solution (10μM) incubated in presence or absence of 20μM EGCG monitored by recording the absorbance at 450nm. (b) Protein content in the unfractionated protein sample (total, T), the supernatant (S) and resuspended pellet (P) after centrifugation of 10μM of refolded TDP-43 samples aggregated in presence or absence of 20μM EGCG was analysed by spotting on a PVDF membrane and staining with Coomassie solution. (c) Representative fluorescence microscopy images of Alexa Fluor Maleimide-labelled TDP-43 incubated with or without EGCG (protein: EGCG ratio 1:2) at 37°C. Scale bar is 25 µm.

Additionally, the particulate nature of the refolded TDP-43 samples incubated with or without EGCG were also investigated *via* a sedimentation assay by fractionating into pellet and supernatant to examine the effect of EGCG on the TDP-43 aggregation [27,46]. For the TDP-43 samples incubated alone, majority of the protein was found in the pellet fraction with only minute quantities present in the supernatant fraction **(Figure 2b)**. In contrast, for TDP-43 co-incubated with EGCG, the pellet fraction did not show any significant protein content and the majority of the protein was found in the supernatant fraction **(Figure 2b)**. This observation further supports an anti-aggregation effect of EGCG on the TDP-43 aggregation *in vitro*. Furthermore, under the mild denaturing conditions (2.5 M urea), when 100 µM TDP-43 was incubated with or without 200 µM EGCG in the aggregation buffer similar anti-aggregation effects of EGCG were observed **(Supplementary Figure S2b)**.

### 3.2. Alexa Fluor-labelled TDP-43 exhibits no aggregate formation in the presence of EGCG in fluorescence microscopy

The effect of EGCG on the formation of TDP-43 aggregates *in vitro* was also analysed using single molecule fluorescence labelling of the TDP-43 protein with Alexa Fluor Malemide-488 [17,24]. To detect the formation of any aggregated TDP-43 species, Alexa Fluor-labelled TDP-43 incubated with or without EGCG were examined under fluorescence microscope using the GFP filter. The TDP-43 samples incubated for 1 h exhibited a few irregular-shaped species, and the number and intensities of such condensed irregular-shaped species further increased with time when visualized up to 12 h post-incubation **(Figure 2c)**. In contrast, the TDP-43 samples incubated in the presence of EGCG did not manifest aggregate formation even when examined at 12 h post-incubation thereby indicating an efficient inhibition of the TDP-43 aggregate formation by EGCG **(Figure 2c)**.

### 3.3. TDP-43 co-incubated with EGCG manifests only small oligomers in AFM imaging

To further visualize the size and morphology, the TDP-43 samples incubated with or without EGCG were visualized using atomic force microscopy (AFM). The TDP-43 sample pre-incubated for 1h in the absence of EGCG when visualized using AFM, showed numerous elongated, irregular shaped aggregates with heights up to ∼ 225 nm (**Figure 3a, 3b** and **3c**). In contrast, the TDP-43 samples pre-incubated with EGCG showed extremely smaller, dispersed, granular structures with heights below ∼ 20 nm **(Figure 3d, 3e** and **3f)**. This suggests that TDP-43 can only form small oligomers in the presence of EGCG, which fail to convert into aggregates thereby supporting an anti-aggregation effect of EGCG on TDP-43. Notably, previously when cultured mammalian cells expressing TDP-43 were treated with EGCG, only oligomer formation, but not aggregate formation, was also observed although the details of the underlying mechanisms or any direct physical interaction of EGCG with TDP-43, were not investigated [21].

**Figure 3:**
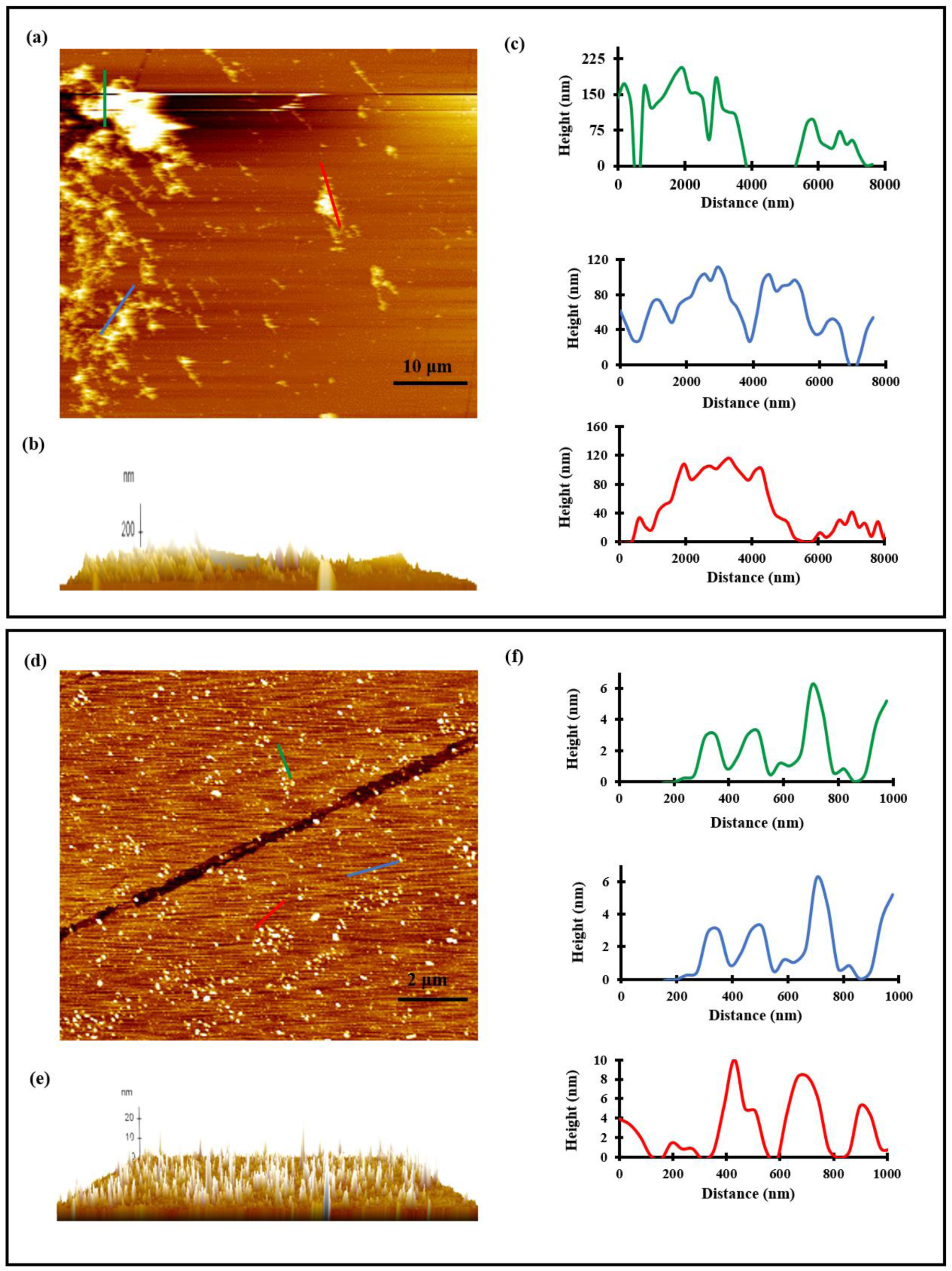
AFM imaging of TDP-43 incubated *in vitro* with or without EGCG. Atomic force microscopy (AFM) images of 100µM TDP-43 aggregates formed in the absence or presence of EGCG (200µM) at 1h post incubation at 37°C. Panels (**a**) and (**d**) represent the AFM images respectively of the samples pre-incubated without and with EGCG. Panels (**b**) and (**e**) show 3D tomographs of the whole image, and panels **(c)** and **(f)** show height-distance profile of the selected oligomers.

### 3.4. EGCG detection with TTC suggests its physical binding with TDP-43 in vitro

Quinones and quinoid-related molecules, such as EGCG, can catalyse redox cycling in the presence of excess glycine as a reductant at an alkaline pH [47]. Tetrazolium salts which are colourless or faintly yellow colour compounds are converted upon reduction into deep coloured compounds known as formazans [48]. EGCG has been shown to reduce tetrazolium dyes to form a coloured product allowing for the detection of EGCG bound to proteins [29,30]. To determine whether EGCG can directly associate with TDP-43, we used a tetrazolium dye, triphenyl tetrazolium chloride (TTC), in a staining assay to detect for protein-bound EGCG molecules. Notably, the TDP-43 samples (100 µM) pre-incubated with EGCG (100 µM) and subjected to extensive dialysis, manifested positive reaction with TTC to generate the coloured formazan product that could be detected by spectrophotometry (500 nm) as well as when visualized by spotting on a PVDF membrane (**Figure 4**). In controls for comparison, when only EGCG (100 µM) or only TDP-43 (100 µM) lacking any EGCG were dialyzed simultaneously and examined similarly by the TTC assay, no coloured product was observed indicating a lack of the EGCG presence at detectable levels (**Figure 4**). This also confirms that the free EGCG could not be retained by the dialysis membrane. Furthermore, the dialysis buffer from the final stage of the dialysis could also not give positive TTC reaction thereby suggesting a lack of presence of detectable EGCG (**Figure 4**). Taken together, the data supports a physical binding of EGCG with TDP-43 that thwarts its removal even upon extensive dialysis.

**Figure 4:**
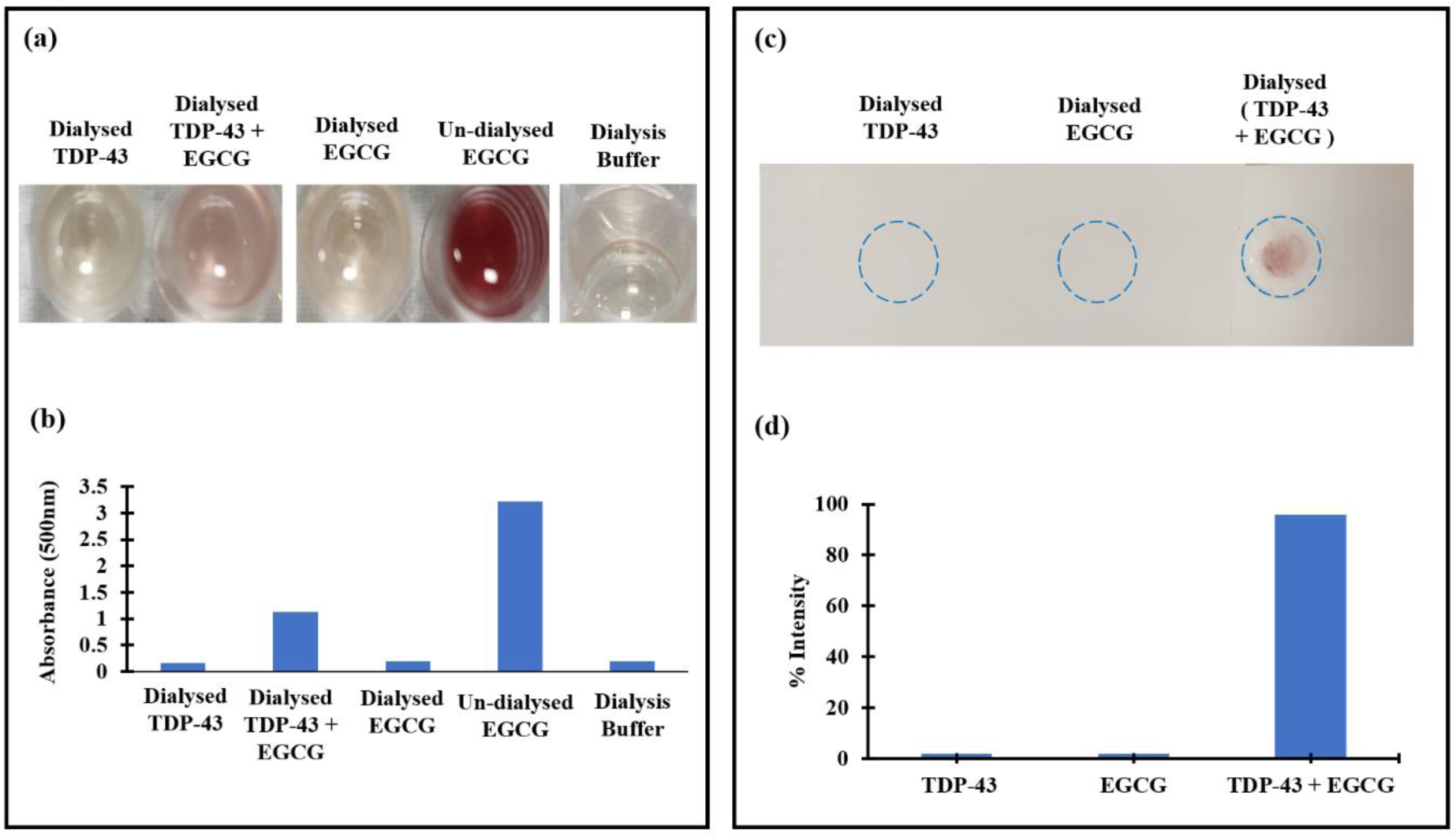
TTC/Glycinate staining of EGCG to examine its *in vitro* binding to TDP-43. **(a)** Reactivity of EGCG bound TDP-43 protein in TTC/glycinate assay in microplate wells. 100μM TDP-43 incubated with 100µM EGCG for 1h and dialysed thoroughly against PBS was subjected to EGCG-specific staining by treating with TTC. Colored product formed upon rection of TTC with EGCG bound to protein was quantified spectrophotometrically. Controls were assayed simultaneously for comparison. **(b)** Reaction of EGCG bound to the TDP-43 protein in TTC/glycinate assay on a PVDF membrane. TDP-43-bound EGCG was detected by first spotting the protein samples dialyzed as in panel (**a**) followed by TTC staining and then quantification by densitometry.

### 3.5. ITC measurements suggest stable binding of TDP-43 with EGCG

The ability of physical binding of EGCG to TDP-43 was also investigated by isothermal titration calorimetry (ITC). To prevent any effect of urea on the molecular interactions between TDP-43 and EGCG, TDP-43 in 6 M urea was diluted 20 times into a refolding buffer with 0.3 M urea in 10 mM Tris (pH 7.5). EGCG was also maintained in 0.3 M urea in 10 mM Tris (pH 7.5) for the titration. The profile for TDP-43 and EGCG titration corrected for dilution effects of EGCG is shown in **Figure 5**. The ITC data obtained for the TDP-43 and EGCG interaction was best fitted using a sequential three-site binding model that indicates one high affinity binding site and two low affinity binding sites. A K_d_ of 7.8 µM was obtained for the high-affinity binding site while the second and third binding sites manifest weaker interactions with K_d_ of 1.4mM and 42.5mM respectively. The enthalpy changes for the binding of EGCG to all three binding sites were negative **(Table 1)** which indicates that the binding of EGCG to TDP-43 is an exothermic process. Notably, the binding of EGCG is a spontaneous process as indicated by the negative free energy changes observed for all the three binding sites **(Table 1)**. The change in the entropy for interaction of EGCG with the first high-affinity binding site of TDP-43 is positive thereby indicating that the binding of EGCG to the high-affinity site is entropy-driven. In contrast, the binding of EGCG to the second and third lower affinity sites was found to be mainly enthalpy-driven as indicated by the negative values of the changes in the entropies **(Table 1)**.

**Figure 5:**
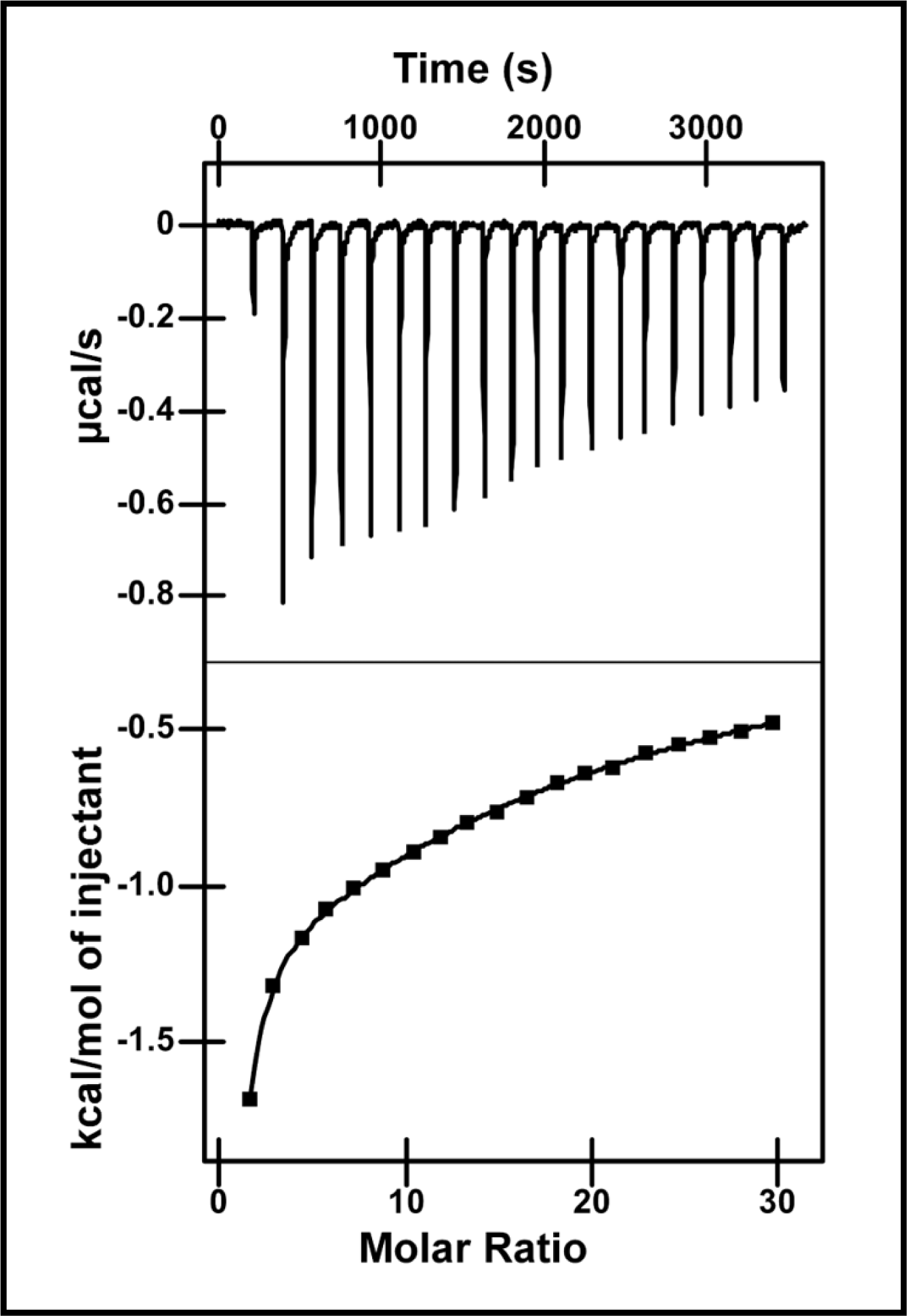
ITC measurements for binding of EGCG to full length TDP-43. **(a)** ITC binding isotherm displaying the raw data obtained from titration of EGCG and TDP-43 at 298K corrected appropriately for heat of dilution. **(b)** Plot representing the amount of heat evolved per mole of EGCG injected against the molar ratio of EGCG to full length TDP-43.

**Table 1:**
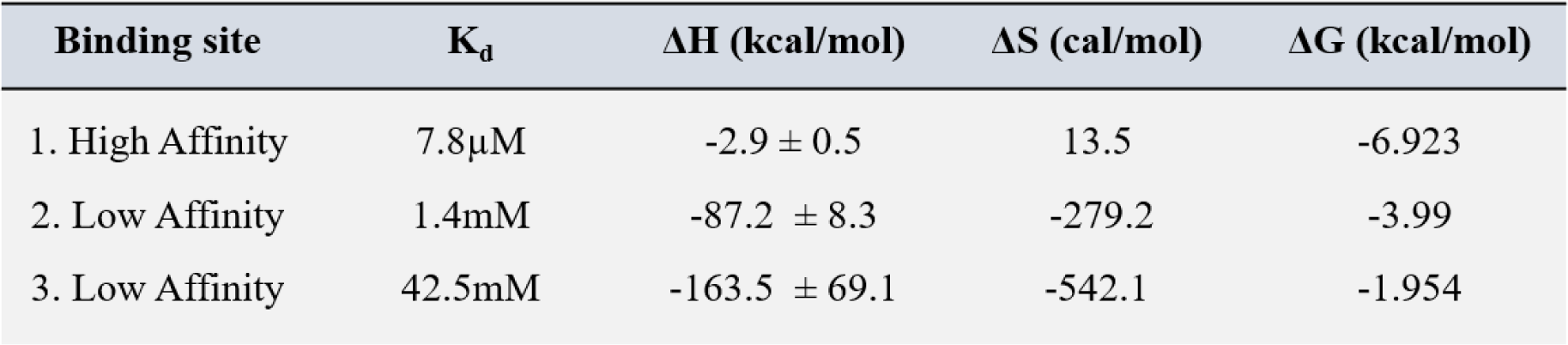
Thermodynamic parameters for EGCG-TDP-43 interactions from isothermal calorimetry (ITC) experiment.

### 3.6. TDP-43 oligomers formed in vitro in presence of EGCG are non-cytotoxic

To analyse any effect of EGCG on the *in vitro* attainment of the cytotoxic conformation by TDP-43, HEK293 cells were treated for 72 h with samples containing either 0.5 µM TDP-43 aggregates formed alone or 0.5 µM TDP-43 oligomers formed in the presence of EGCG (protein: EGCG - 1:2). When an MTT cell viability assay was used to assess the cytotoxicity of these TDP-43 aggregates/oligomers, the HEK293 cells treated with TDP-43 aggregates showed a significant decrease in the cell viability (87.75%) in comparison with the untreated HEK293 cells (p < 0.0001) whereas, in contrast, the TDP-43 oligomers formed in the presence of EGCG did not cause decrease in the cell viability and rather even significantly increased the viability of the cells to 116.9% (p < 0.001). Although the reason for this enhancement of the cell viability is not understood, the data overall, however, support that while the TDP-43 aggregates formed *in vitro* can cause cytotoxicity, the oligomers of TDP-43 formed *in vitro* in the presence of EGCG are not cytotoxic **(Figure 6)**.

**Figure 6:**
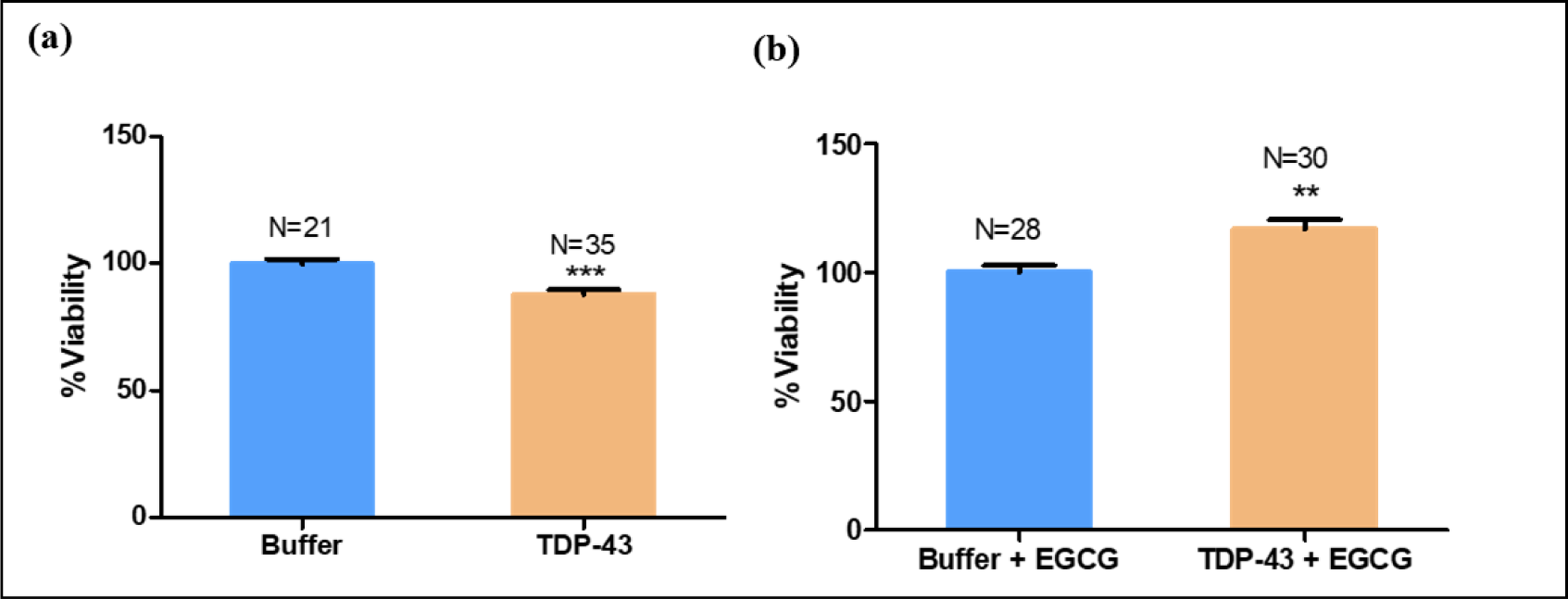
MTT cell viability assay of HEK293 cells treated with TDP-43 oligomers formed *in vitro* with EGCG. Cytotoxicity assessment of TDP-43 aggregates/oligomers prepared in the absence or presence of EGCG. (**a**) Percent viability plotted upon treatment of HEK293 with 0.5 µM TDP-43 aggregates formed in absence of EGCG. (**b**) Percent viability plotted upon treatment of HEK-293 with 0.5 µM TDP-43 oligomers formed in the presence of EGCG. N = number of wells analysed for each sample. *** represents *p* ≤ 0.0001. **represents *p* ≤ 0.001. *p*-value represents the significance of the difference between the respective buffer and the treated samples.

### 3.7. In silico molecular docking predicts binding sites of EGCG on TDP-43

As EGCG showed anti-TDP-43 aggregation propensity and exhibited physical binding capability with TDP-43, we further used *in silico* molecular docking tool, AutoDock 4.2, to predict binding site(s) of EGCG on the TPD-43 protein. So far, the structure of the full-length TDP-43 protein is not available, but the structures of certain domains and fragments of TDP-43 protein have been elucidated [31–33,49,50].

When we performed *in silico* blind docking of the ligand, EGCG, with different domain structures of TDP-43 reported in the Protein Data Bank (PDB), the best docking score (-11.31 kcal/mol) and inhibitory constant (log Ki: -2.25; Ki: 5.15 nM) were observed for the structure of the low complexity domain of TDP-43 (aa: 276-414; PDB ID: 7KWZ) which is a cryo-EM structure of the oligomeric ensemble of this domain in an amyloid-like conformation [33]. This structure contains 14 β-strands where the N-terminal of this protein fragment has hydrophobic residues stabilizing the region and the C-terminal region is stabilized by steric-zipper interactions within the strands β9, β10, β11 and β13 [33]. AutoDock 4.2 predicted EGCG to dock to the relatively hydrophobic and non-planar N-terminal region of the low complexity domain of TDP-43 involving residues Phe-313 (Chain E), Ala-341 (Chain D) that are within the strong hydrophobic core stabilizing the region.

Additionally, AutoDock 4.2 also predicted appreciable and relatively comparable docking scores (∼ -10 kcal/mol) of EGCG to the crystal structure of the N-terminal domain of TDP-43 (aa:1-80; PDB ID: 5MDI) and the NMR structure of the RRM1-2 domains of TDP-43 (aa: 96-269; PDB ID: 4BS2) irrespective of whether the RNA recognition motifs (RRMs) were bound or unbound to RNA [31,32] (**Figure 7a** and **7b**). Furthermore, an appreciable docking scores (∼ -9.7 kcal/mol) of EGCG was also predicted to another structure of the N-terminal domain of TDP-43 (aa:1-77; PDB ID: 2N4P) suggesting a possibility of binding (**Figure 7a** and **7b**). The docking of EGCG to several other available structures of the domains and peptides of TDP-43 did not show appreciable docking scores thereby suggesting a lack of predicted binding of EGCG with these structures (**Figure 7a** and **7b**).

**Figure 7:**
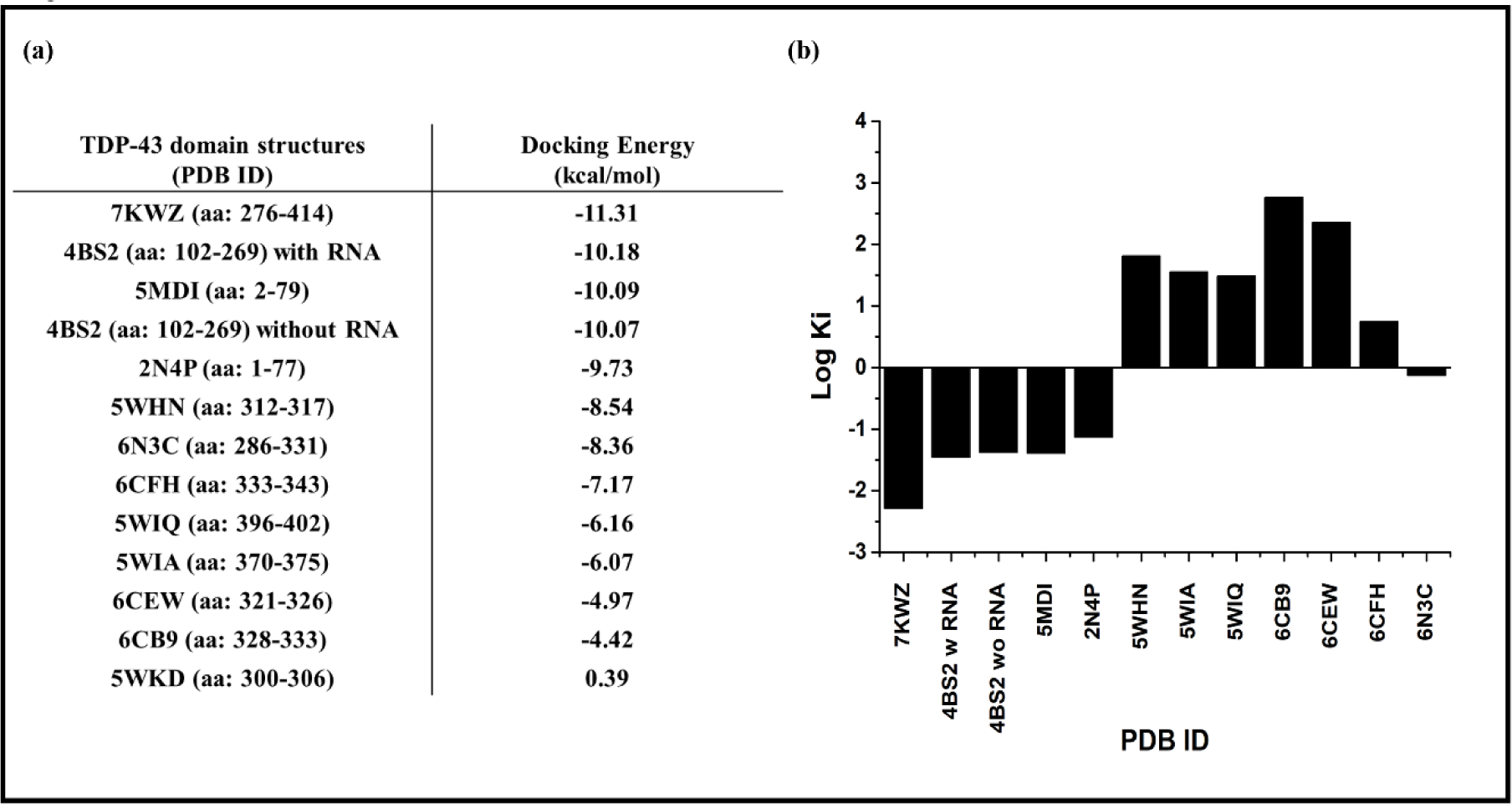
*In silico* molecular docking studies of the structures of domains or fragments of TDP-43 with EGCG. **(a)** Molecular docking scores of the domain structures or fragments of TDP-43 with EGCG. The PDB ID of the various domains or fragments of TDP-43 along with the amino acids comprising the domains or fragments are mentioned along with their individual docking score after blind docking with AutoDock 4.2. **(b)** The column chart of the log K_i_ values of the binding of EGCG with the individual domain structures or the fragments of TDP-43 as yielded by AutoDock 4.2.

### 3.8. Molecular dynamics simulations support stable interaction of EGCG with the aggregation-prone C-terminal domain of TDP-43

To study the *in-situ* structure, binding and dynamics of the EGCG bound complex structure, the ensemble structure of the low complexity C-terminal domain (CTD) of TDP-43 (aa: 276-414; PDB ID: 7KWZ) with EGCG, which showed best docking score using AutoDock 4.2, was simulated using all-atom MD simulations. The RMSD of protein-ligand complex was observed to converge and stabilize when MD simulation was carried for 300 ns (**Supplementary Figures S3** and **S4**). Furthermore, during the whole free equilibration MD simulation of 300 ns, ECGG remained bound to the structure of the CTD of TDP-43 **(Figure 8a)**. In contrast, EGCG complexed with the N-terminal domain structure (PDB ID: 5MDI) or the RRM1-2 domain structure (PDB ID: 4BS2) showed discontinuous contacts even when MD simulations were performed to an extended time of 800 ns **(Figure 8b** and **Supplementary Figures S3** and **S4)**. The MM-PBSA calculations for the free energy of binding of EGCG employing the gmx_MMPBSA tool and using the last 25 ns MD simulation data yielded -2.04 kcal/mol, -7.94 kcal/mol and -20.29 kcal/mol binding energies respectively with the N-terminal domain, RRM1-2 domain and CTD structures thereby suggesting a relatively more stable binding of EGCG to the CTD as compared to the other two domains **(Figure 8c, 8d** and **Supplementary Table S1)**. Hence, further analysis of the types of bonds/contacts that stabilise the interaction of EGCG to CTD was performed using the tool, PyContact. This analysis showed that the residues Ala-341 of chain C and D, Ser-342 of chain D and E, Phe-313, Phe-316 and Gln-343 of chain E of the ensemble structure of the CTD contributed significantly towards the interaction with EGCG through H-bonding and other non-covalent interactions **(Figure 8e)**. Furthermore, a clustering analysis gave a single large cluster followed by a cluster with a unary structure for the 300 ns simulation of the CTD complexed with EGCG and the representative structure from the large cluster showed H-bond interactions between the hydroxyl groups of EGCG and the residues Ser-342, Gln-343 of the chain E, Ala-341 and Ser-342 of the chain D and Ala-341 of chain C of the ensemble structure of CTD. Additionally, other non-covalent interactions including a pi-pi interaction between the benzopyran ring of EGCG and Phe-316 from chain E and the side benzene ring of EGCG with Phe-313 from chain E were also observed. Also a pi-sigma interaction was predicted between EGCG and Ala-341 from chain E of the oligomeric CTD structure **(Figure 8f** and **Supplementary Movie S1)**.

**Figure 8:**
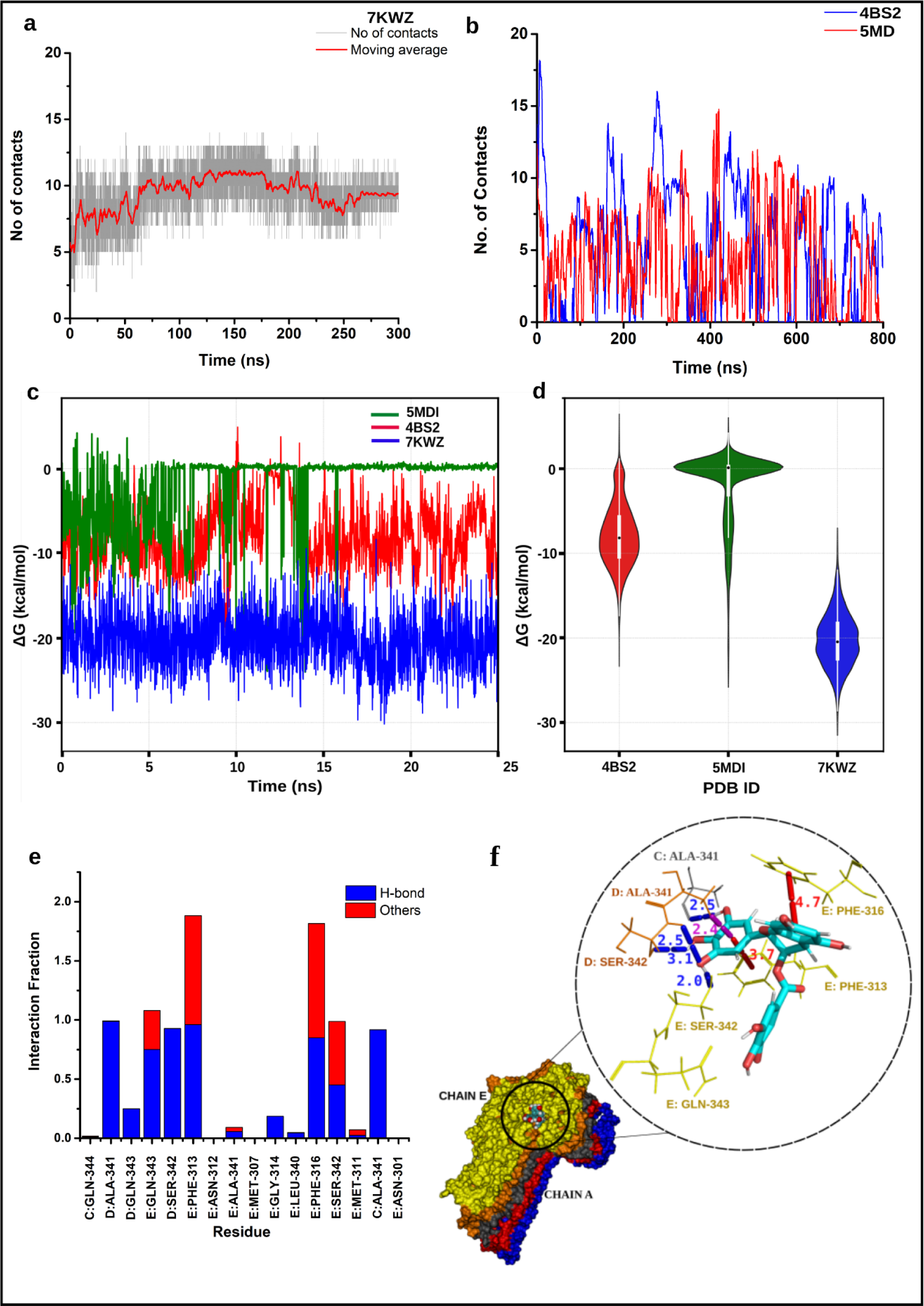
All-atom MD simulation results of domains of TDP-43 complexed with EGCG. **(a)** Number of contacts formed between 7KWZ with EGCG for the 300 ns simulation (grey) along with the moving average of the number of contacts (red). **(b)** The moving average of the number of contacts formed between 5MDI (red) and 4BS2 (blue) with EGCG during the 800 ns simulation. **(c)** The total free energy of binding of the domains of TDP-43 with EGCG for the last 25 ns simulation. **(d)** The violin plot of the distribution of the values of the free energy of binding for the last 25 ns of the three domains of TDP-43 with EGCG. The box plot inside each violin shows the interquartile ranges along with the median value marked as white circle. The ensemble structure of the C-terminal domain of TDP-43 (PDB ID: 7KWZ) showed the highest negative free energy of binding as compared to the other two domains. **(e)** The stacked column chart representing the interacting residues of the C-terminal of TDP-43 with EGCG also showing the individual contribution of the different type of interactions such as hydrogen (blue) and other non-covalent interactions (red) with EGCG. The interaction fraction is the measure of the lifetime of the individual interactions during the simulation time. The value of 1 means the interaction is present throughout the simulation time. Residues that are predicted to be interacting equally through both hydrogen and other non-covalent interaction for most frames reached values above 1. **(f)** The representative structure of 7KWZ in complex with the ligand from the clustering performed for 300 ns simulation with a cut-off of 2Å is shown and the close view of the type of interactions predicted between the residues of the protein and EGCG with their distances labelled in Angstrom units. Red – Pi-Pi interactions, Blue – Hydrogen bond; Purple - Pi-sigma interaction.

## 4. Discussion

Various small molecules of natural origin such as curcumin, EGCG, resveratrol, catechin etc. have been demonstrated to efficiently inhibit the formation of certain amyloid fibrils and their related cytotoxicity [51]. As the polyphenol epigallocatechine gallate (EGCG) is a natural molecule being a component of a popular beverage, green tea, we have examined its inhibitory potential towards the *in vitro* aggregation of TDP-43 protein which is associated with ALS and FTLD-TDP diseases. Previous studies have also reported that EGCG can effectively modulate the misfolding and aggregation of certain amyloidogenic proteins like α-synuclein, amyloid-β, huntingtin and tau [19,30,52,53].

In this study, we illustrate the binding and the inhibitory potential of EGCG on full-length TDP-43 by *in silico* and *in vitro* biochemical assays. Solution turbidity measurements, sedimentation assay along with microscopic visualization of Alexa-fluor labelled fluorescent TDP-43 aggregates in the presence of EGCG revealed that EGCG can thwart the TDP-43’s aggregate formation *in vitro* [17,24]. EGCG has been previously shown to bind unfolded polypeptides preventing their fibril formation and resulting in seed-incompetent and non-toxic off-pathway oligomeric structures [19,30,54]. Similarly, our AFM imaging suggests that EGCG diminishes the formation of otherwise irregularly shaped filament-like aggregates of TDP-43 and exhibits only small spheroidal oligomers. Thus, we demonstrate that EGCG might bind to TDP-43 arresting the protein in the oligomeric stage and further disallowing to its follow its fibrillation pathway.

In a previous study using HEK293T cells, EGCG prevented the degradation of functional TDP-43 and promoted the formation of functional non-cytotoxic oligomers in the nuclei and EGCG was proposed to exert this effect by stabilizing the C-terminal domain of TDP-43 however, any physical binding of EGCG to TDP-43 was not examined [21]. Therefore, here we explored the physical binding of EGCG with TDP-43 by employing several methods such as tetrazolium staining, isothermal calorimetry and *in silico* molecular docking and molecular dynamic simulation (MDS) studies. Previously, specific detection of EGCG-bound proteins by staining with tetrazolium dyes has been utilised for studying the direct binding of EGCG to proteins [19,53]. Our triphenyl tetrazolium (TTC) staining of dialysed TDP-43 aggregates formed in the presence of EGCG manifested the formation of a coloured formazan product indicating that EGCG physically binds with TDP-43. Furthermore, the ITC experiment for examining the EGCG interaction with TDP-43 also suggests that TDP-43 presents multiple binding sites for EGCG with at least one high-affinity and two low-affinity interaction sites. Notably, similarl multiple binding events-strong initial binding followed by a secondary weaker binding event have been reported for EGCG with tau and HSA proteins [52,55]. Thermodynamic parameters obtained from the ITC study revealed that the high-affinity binding of EGCG to TDP-43 is entropy-driven while the binding to low-affinity TDP-43 sites is enthalpy-driven. Notably, previous studies have also reported the similar synergistic action of entropic and enthalpy forces driving the binding interactions between BSA and various ligands [56,57].

To examine the relative cytotoxicity of the TDP-43 aggregates formed in the presence or absence of EGCG, we assessed the toxicity of these aggregates/oligomers towards HEK cells *via* MTT reduction assay. While the addition of TDP-43 aggregates pre-formed *in vitro* in absence of EGCG was mildly toxic to the HEK cells, the TDP-43 oligomers formed consequent upon its pre-incubation with EGCG, did not exert cytotoxicity. This observation complements the previous observations where EGCG induced the formation of non-toxic functional oligomers of TDP-43 in nuclei of HEK293T cells endogenously expressing TDP-43. Similar effect of EGCG has also been demonstrated with tau protein where toxicity conferred by exogenous tau aggregates was significantly reduced when cells were treated with tau species formed in the presence of EGCG [19,53].

As our *in vitro* experiments revealed a physical interaction between full length TDP-43 and EGCG, to obtain further insights into the mechanisms and potential binding sites, we performed *in-silico* molecular docking and molecular dynamic simulations (MDS) studies. To date, the globular structure of full-length TDP-43 is not elucidated thus we have performed the *in silico* studies with the available domain structures of TDP-43. Previous studies have shown that small molecules can bind to the N-terminal of TDP-43 allosterically modulating the RNA binding of the protein and rescuing it from locomotor defects in the *Drosophila* model of ALS [10]. Similarly, various small molecules have been screened and studied for their interaction directly with the RRM1-2 domains [12]. Our simulation results of the apo-structure of the RRM1-2 domains with EGCG showed different binding regions during the 800 ns simulation run which might be the result of the dynamic conformation changes and exposure of certain buried residues during the simulation. Our results are consistent with the previous reports showing the dynamic conformation changes during the MD simulation run for the apo-structure of tandem RRM1-2 domains [58]. However, EGCG remained stably bound to the hydrophobic, non-planar, amino terminal region of the ensemble structure of the low complexity C-terminal domain of TDP-43 (PDB ID: 7KWZ, aa: 276-414) for the 300 ns simulation performed, unlike during the simulation performed with both the N-terminal (PDB ID: 5MDI, aa: 1-80) and the RRM1-2 domain (PDB ID: 4BS2, aa: 96-269). The interaction between Phe-313, Ala-341 with Ile-318, Met-336 residues are due to non-planar conformation of the N-terminal region of the LCD, thereby forming rugged surfaces exposing hydrophobic residues that might facilitate the recruitment of TDP-43 monomers to form amyloid fibrils [46]. Interestingly, our study predicted the binding of EGCG with the Phe-313 and Ala-341 residues of the LCD structure thereby explaining the possibility of interaction of EGCG with the exposed hydrophobic pocket residues and preventing the fibril formation. Moreover, the predicted region of interaction has no reported history of carrying a mutation except a silent mutation Q343R and no further conformational changes are reported for different mechanisms of action [33]. The docking of EGCG with the apo form or RNA bound form of TDP-43 showed interaction near RRM2 domain whereas 800 ns simulation data showed that EGCG bound transiently to different regions in 4BS2 RRM1-2 structure and 750-800s period of MDS exhibited the interaction near RRM1, that is reported to be essential but not sufficient for stable interaction with the RNA and generating multimers [59]. The N-terminal region of TDP-43 is required for the formation of different oligomers that are functional and sterically hinder the aggregation mediated by CTD of different TDP-43 monomers [31]. Hence, unstable interactions of EGCG with the RRM12 domains of TDP-43 from our simulation study can be an indication of the effect of the small molecule on the aggregation-prone domains (LCD) of TDP-43 and not on the domains involved in the functional oligomer formation, which needs further validation through interaction studies with different domains of TDP-43.

Overall, this study shows that EGCG binds to TDP-43 and impedes its aggregation and allows for the formation of only small oligomeric TDP-43 species that are non-cytotoxic, making EGCG a possible candidate molecule for potential therapeutic research for TDP-43 proteinopathies like ALS.

## 5. Conclusion

Here, we report that the *in vitro* aggregation of full length TDP-43 is inhibited by the green tea polyphenol, EGCG, resulting in the formation of only small oligomers. A physical binding of EGCG with TDP-43 was also demonstrated. Furthermore, ITC binding studies also suggest the presence of multiple possible binding sites for EGCG on TDP-43. We find that the *in vitro*-made aggregates of TDP-43 can cause mild cytotoxicity when added to the HEK293 cell culture media whereas, strikingly, the small oligomers of TDP-43 formed in the presence of EGCG are not cytotoxic. Additionally, *in silico* molecular docking and MD simulation studies predict stable interaction of EGCG with the amyloid-like ensemble structure formed by the low complexity C-terminal region of TDP-43. Taken together, the data supports that the green tea polyphenol, EGCG, has *in vitro* anti-aggregation propensity against TDP-43 thereby making it non-cytotoxic. These findings would be relevant to the ongoing quest for therapeutic molecules against the TDP-43 aggregation, the extracellular TDP-43’s prion-like propagation and the TDP-43 proteinopathies in general.

## Acknowledgements

We thank IIT Hyderabad, funded by Ministry of Education (MoE), Govt. of India, for research infrastructure and support. VDM is thankful to UGC, Govt. of India, for junior research fellowship (JRF). PS thanks MoE, Govt. of India, for senior research fellowship (SRF). YS thanks CSIR, Govt. of India, for SRF. RB thanks MoE, Govt. of India, for Prime Minister Research Fellowship (PMRF). AB thanks MoE, Govt. of India, for senior research fellowship (SRF). We thank Yoshiaki Furukawa, Keio University, for a plasmid gift. We thank Sri Chengappa Thumisi from IIT-Hyderabad for help with AFM imaging. Basant K Patel thanks SERB-DST, Govt. of India for a research grant (Grant no: SERB/CRG/2021/006856).

## References

[1] A. Prasad, V. Bharathi, V. Sivalingam, A. Girdhar, B.K. Patel, Molecular Mechanisms of TDP-43 Misfolding and Pathology in Amyotrophic Lateral Sclerosis, Front. Mol. Neurosci. 12 (2019). https://www.frontiersin.org/articles/10.3389/fnmol.2019.00025.

[2] L. François-Moutal, S. Perez-Miller, D.D. Scott, V.G. Miranda, N. Mollasalehi, M. Khanna, Structural Insights Into TDP-43 and Effects of Post-translational Modifications, Front. Mol. Neurosci. 12 (2019) 301. 10.3389/fnmol.2019.00301.

[3] S. Maharana, J. Wang, D.K. Papadopoulos, D. Richter, A. Pozniakovsky, I. Poser, M. Bickle, S. Rizk, J. Guillén-Boixet, T.M. Franzmann, M. Jahnel, L. Marrone, Y.-T. Chang, J. Sterneckert, P. Tomancak, A.A. Hyman, S. Alberti, RNA buffers the phase separation behavior of prion-like RNA binding proteins, Science. 360 (2018) 918–921. 10.1126/science.aar7366.

[4] K. Matsukawa, M.S. Kukharsky, S.-K. Park, S. Park, N. Watanabe, T. Iwatsubo, T. Hashimoto, S.W. Liebman, T.A. Shelkovnikova, Long non-coding RNA NEAT1_1 ameliorates TDP-43 toxicity in in vivo models of TDP-43 proteinopathy, RNA Biol. 18 1546–1554. 10.1080/15476286.2020.1860580.

[5] K.C. Luk, V.M. Kehm, B. Zhang, P. O’Brien, J.Q. Trojanowski, V.M.Y. Lee, Intracerebral inoculation of pathological α-synuclein initiates a rapidly progressive neurodegenerative α-synucleinopathy in mice, J. Exp. Med. 209 (2012) 975–986. 10.1084/jem.20112457.

[6] T. Kanouchi, T. Ohkubo, T. Yokota, Can regional spreading of amyotrophic lateral sclerosis motor symptoms be explained by prion-like propagation, J. Neurol. Neurosurg. Psychiatry. 83 (2012) 739–745. 10.1136/jnnp-2011-301826.

[7] S. Porta, Y. Xu, C.R. Restrepo, L.K. Kwong, B. Zhang, H.J. Brown, E.B. Lee, J.Q. Trojanowski, V.M.-Y. Lee, Patient-derived frontotemporal lobar degeneration brain extracts induce formation and spreading of TDP-43 pathology in vivo, Nat. Commun. 9 (2018) 4220. 10.1038/s41467-018-06548-9.

[8] M.S. Feiler, B. Strobel, A. Freischmidt, A.M. Helferich, J. Kappel, B.M. Brewer, D. Li, D.R. Thal, P. Walther, A.C. Ludolph, K.M. Danzer, J.H. Weishaupt, TDP-43 is intercellularly transmitted across axon terminals, J. Cell Biol. 211 (2015) 897–911. 10.1083/jcb.201504057.

[9] S.T. Kumar, S. Nazarov, S. Porta, N. Maharjan, U. Cendrowska, M. Kabani, F. Finamore, Y. Xu, V.M.-Y. Lee, H.A. Lashuel, Seeding the aggregation of TDP-43 requires post-fibrillization proteolytic cleavage, Nat. Neurosci. 26 (2023) 983–996. 10.1038/s41593-023-01341-4.

[10] N. Mollasalehi, L. Francois-Moutal, D.D. Scott, J.A. Tello, H. Williams, B. Mahoney, J.M. Carlson, Y. Dong, X. Li, V.G. Miranda, V. Gokhale, W. Wang, S.J. Barmada, M. Khanna, An Allosteric Modulator of RNA Binding Targeting the N-Terminal Domain of TDP-43 Yields Neuroprotective Properties, ACS Chem. Biol. 15 (2020) 2854–2859. 10.1021/acschembio.0c00494.

[11] L. McGurk, E. Gomes, L. Guo, J. Shorter, N.M. Bonini, Poly(ADP-ribose) Engages the TDP-43 Nuclear-Localization Sequence to Regulate Granulo-Filamentous Aggregation, Biochemistry. 57 (2018) 6923–6926. 10.1021/acs.biochem.8b00910.

[12] M.Y. Fang, S. Markmiller, A.Q. Vu, A. Javaherian, W.E. Dowdle, P. Jolivet, P.J. Bushway, N.A. Castello, A. Baral, M.Y. Chan, J.W. Linsley, D. Linsley, M. Mercola, S. Finkbeiner, E. Lecuyer, J.W. Lewcock, G.W. Yeo, Small-Molecule Modulation of TDP-43 Recruitment to Stress Granules Prevents Persistent TDP-43 Accumulation in ALS/FTD, Neuron. 103 (2019) 802–819.e11. 10.1016/j.neuron.2019.05.048.

[13] D.F. Tardiff, M.L. Tucci, K.A. Caldwell, G.A. Caldwell, S. Lindquist, Different 8-Hydroxyquinolines Protect Models of TDP-43 Protein, α-Synuclein, and Polyglutamine Proteotoxicity through Distinct Mechanisms, J. Biol. Chem. 287 (2012) 4107–4120. 10.1074/jbc.M111.308668.

[14] X. Liu, S. Duan, Y. Jin, E. Walker, M. Tsao, J.H. Jang, Z. Chen, A.K. Singh, K.L. Cantrell, H.I. Ingolfsson, S.K. Buratto, M.T. Bowers, Computationally Designed Molecules Modulate ALS-Related Amyloidogenic TDP-43307–319 Aggregation, ACS Chem. Neurosci. (2023). 10.1021/acschemneuro.3c00582.

[15] M.R. Salaikumaran, P.P. Gopal, Rational Design of TDP-43 Derived α-Helical Peptide Inhibitors: an In-Silico Strategy to Prevent TDP-43 Aggregation in Neurodegenerative Disorders, (2023) 2023.10.26.564235. 10.1101/2023.10.26.564235.

[16] A. Prasad, G. Raju, V. Sivalingam, A. Girdhar, M. Verma, A. Vats, V. Taneja, G. Prabusankar, B.K. Patel, An acridine derivative, [4,5-bis{(N-carboxy methyl imidazolium)methyl}acridine] dibromide, shows anti-TDP-43 aggregation effect in ALS disease models, Sci. Rep. 6 (2016) 39490. 10.1038/srep39490.

[17] A. Girdhar, V. Bharathi, V.R. Tiwari, S. Abhishek, W. Deeksha, U.S. Mahawar, G. Raju, S.K. Singh, G. Prabusankar, E. Rajakumara, B.K. Patel, Computational insights into mechanism of AIM4-mediated inhibition of aggregation of TDP-43 protein implicated in ALS and evidence for in vitro inhibition of liquid-liquid phase separation (LLPS) of TDP-432C-A315T by AIM4, Int. J. Biol. Macromol. 147 (2020) 117–130. 10.1016/j.ijbiomac.2020.01.032.

[18] W. Dai, C. Ruan, Y. Zhang, J. Wang, J. Han, Z. Shao, Y. Sun, J. Liang, Bioavailability enhancement of EGCG by structural modification and nano-delivery: A review, J. Funct. Foods. 65 (2020) 103732. 10.1016/j.jff.2019.103732.

[19] D.E. Ehrnhoefer, J. Bieschke, A. Boeddrich, M. Herbst, L. Masino, R. Lurz, S. Engemann, A. Pastore, E.E. Wanker, EGCG redirects amyloidogenic polypeptides into unstructured, off-pathway oligomers, Nat. Struct. Mol. Biol. 15 (2008) 558–566. 10.1038/nsmb.1437.

[20] S.-H. Koh, S.M. Lee, H.Y. Kim, K.-Y. Lee, Y.J. Lee, H.-T. Kim, J. Kim, M.-H. Kim, M.S. Hwang, C. Song, K.-W. Yang, K.W. Lee, S.H. Kim, O.-H. Kim, The effect of epigallocatechin gallate on suppressing disease progression of ALS model mice, Neurosci. Lett. 395 (2006) 103–107. 10.1016/j.neulet.2005.10.056.

[21] I.-F. Wang, H.-Y. Chang, S.-C. Hou, G.-G. Liou, T.-D. Way, C.-K. James Shen, The self-interaction of native TDP-43 C terminus inhibits its degradation and contributes to early proteinopathies, Nat. Commun. 3 (2012) 766. 10.1038/ncomms1766.

[22] Y. Furukawa, K. Kaneko, S. Watanabe, K. Yamanaka, N. Nukina, A Seeding Reaction Recapitulates Intracellular Formation of Sarkosyl-insoluble Transactivation Response Element (TAR) DNA-binding Protein-43 Inclusions, J. Biol. Chem. 286 (2011) 18664–18672. 10.1074/jbc.M111.231209.

[23] S. Nirwal, P. Saravanan, A. Bajpai, V.D. Meshram, G. Raju, W. Deeksha, G. Prabusankar, B.K. Patel, In Vitro Interaction of a C-Terminal Fragment of TDP-43 Protein with Human Serum Albumin Modulates Its Aggregation, J. Phys. Chem. B. 126 (2022) 9137–9151. 10.1021/acs.jpcb.2c04469.

[24] S. Preethi, V. Bharathi, B.K. Patel, Zn2+ modulates in vitro phase separation of TDP-432C and mutant TDP-432C-A315T C-terminal fragments of TDP-43 protein implicated in ALS and FTLD-TDP diseases, Int. J. Biol. Macromol. 176 (2021) 186–200. 10.1016/j.ijbiomac.2021.02.054.

[25] V. Sivalingam, N.L. Prasanna, N. Sharma, A. Prasad, B.K. Patel, Wild-type hen egg white lysozyme aggregation in vitro can form self-seeding amyloid conformational variants, Biophys. Chem. 219 (2016) 28–37. 10.1016/j.bpc.2016.09.009.

[26] V. Sivalingam, B.K. Patel, Familial mutations in fibrinogen Aα (FGA) chain identified in renal amyloidosis increase in vitro amyloidogenicity of FGA fragment, Biochimie. 127 (2016) 44–49. 10.1016/j.biochi.2016.04.020.

[27] A. Prasad, V. Sivalingam, V. Bharathi, A. Girdhar, B.K. Patel, The amyloidogenicity of a C-terminal region of TDP-43 implicated in Amyotrophic Lateral Sclerosis can be affected by anions, acetylation and homodimerization, Biochimie. 150 (2018) 76–87. 10.1016/j.biochi.2018.05.003.

[28] C.T. Rueden, J. Schindelin, M.C. Hiner, B.E. DeZonia, A.E. Walter, E.T. Arena, K.W. Eliceiri, ImageJ2: ImageJ for the next generation of scientific image data, BMC Bioinformatics. 18 (2017) 529. 10.1186/s12859-017-1934-z.

[29] A. Somayaji, C. Shastry, Interference of Antioxidant Flavonoids with MTT Tetrazolium Assay in a Cell Free System, J. Pharm. Res. Int. (2021) 76–83. 10.9734/jpri/2021/v33i49A33305.

[30] J. Bieschke, J. Russ, R.P. Friedrich, D.E. Ehrnhoefer, H. Wobst, K. Neugebauer, E.E. Wanker, EGCG remodels mature alpha-synuclein and amyloid-beta fibrils and reduces cellular toxicity, Proc. Natl. Acad. Sci. U. S. A. 107 (2010) 7710–7715. 10.1073/pnas.0910723107.

[31] T. Afroz, E.M. Hock, P. Ernst, C. Foglieni, M. Jambeau, L.A.B. Gilhespy, F. Laferriere, Z. Maniecka, A. Plückthun, P. Mittl, P. Paganetti, F.H.T. Allain, M. Polymenidou, Functional and dynamic polymerization of the ALS-linked protein TDP-43 antagonizes its pathologic aggregation, Nat Commun. 8 (2017) 45. 10.1038/s41467-017-00062-0.

[32] P.J. Lukavsky, D. Daujotyte, J.R. Tollervey, J. Ule, C. Stuani, E. Buratti, F.E. Baralle, F.F. Damberger, F.H.-T. Allain, Molecular basis of UG-rich RNA recognition by the human splicing factor TDP-43, Nat. Struct. Mol. Biol. 20 (2013) 1443–1449. 10.1038/nsmb.2698.

[33] Q. Li, W.M. Babinchak, W.K. Surewicz, Cryo-EM structure of amyloid fibrils formed by the entire low complexity domain of TDP-43, Nat. Commun. 12 (2021) 1620. 10.1038/s41467-021-21912-y.

[34] Lindahl, Abraham, Hess, V.D. Spoel, GROMACS 2019 Manual, (2018). 10.5281/ZENODO.2424486.

[35] K. Vanommeslaeghe, E. Hatcher, C. Acharya, S. Kundu, S. Zhong, J. Shim, E. Darian, O. Guvench, P. Lopes, I. Vorobyov, A.D. MacKerell, CHARMM General Force Field (CGenFF): A force field for drug-like molecules compatible with the CHARMM all-atom additive biological force fields, J. Comput. Chem. 31 (2010) 671–690. 10.1002/jcc.21367.

[36] W. Yu, X. He, K. Vanommeslaeghe, A.D. MacKerell, Extension of the CHARMM General Force Field to Sulfonyl-Containing Compounds and Its Utility in Biomolecular Simulations, J. Comput. Chem. 33 (2012) 2451–2468. 10.1002/jcc.23067.

[37] T. Darden, D. York, L. Pedersen, Particle mesh Ewald: An N⋅log(N) method for Ewald sums in large systems, J. Chem. Phys. 98 (1993) 10089–10092. 10.1063/1.464397.

[38] H.J.C. Berendsen, J.P.M. Postma, W.F. van Gunsteren, A. DiNola, J.R. Haak, Molecular dynamics with coupling to an external bath, J. Chem. Phys. 81 (1984) 3684–3690. 10.1063/1.448118.

[39] M. Scheurer, P. Rodenkirch, M. Siggel, R.C. Bernardi, K. Schulten, E. Tajkhorshid, T. Rudack, PyContact: Rapid, Customizable, and Visual Analysis of Noncovalent Interactions in MD Simulations, Biophys. J. 114 (2018) 577–583. 10.1016/j.bpj.2017.12.003.

[40] M.S. Valdés-Tresanco, M.E. Valdés-Tresanco, P.A. Valiente, E. Moreno, gmx_MMPBSA: A New Tool to Perform End-State Free Energy Calculations with GROMACS, J. Chem. Theory Comput. 17 (2021) 6281–6291. 10.1021/acs.jctc.1c00645.

[41] B.R.I. Miller, T.D.Jr. McGee, J.M. Swails, N. Homeyer, H. Gohlke, A.E. Roitberg, MMPBSA.py: An Efficient Program for End-State Free Energy Calculations, J. Chem. Theory Comput. 8 (2012) 3314–3321. 10.1021/ct300418h.

[42] W. Humphrey, A. Dalke, K. Schulten, VMD: visual molecular dynamics, J. Mol. Graph. 14 (1996) 33–38, 27–28. 10.1016/0263-7855(96)00018-5.

[43] B.D.S. Visualizer, Dassault Systemes; San Diego, CA, USA: 2021, Version V21. 1 (2021) 20298.

[44] The PyMOL Molecular Graphics System, Version 1.2r3pre, Schrödinger, LLC., (n.d.).

[45] M. Vivoli Vega, A. Nigro, S. Luti, C. Capitini, G. Fani, L. Gonnelli, F. Boscaro, F. Chiti, Isolation and characterization of soluble human full-length TDP-43 associated with neurodegeneration, FASEB J. Off. Publ. Fed. Am. Soc. Exp. Biol. 33 (2019) 10780–10793. 10.1096/fj.201900474R.

[46] C.O. Zlatic, Y. Mao, T.M. Ryan, Y.-F. Mok, B.R. Roberts, G.J. Howlett, M.D.W. Griffin, Fluphenazine·HCl and Epigallocatechin Gallate Modulate the Rate of Formation and Structural Properties of Apolipoprotein C-II Amyloid Fibrils, Biochemistry. 54 (2015) 3831–3838. 10.1021/acs.biochem.5b00399.

[47] M.A. Paz, R. Flückiger, A. Boak, H.M. Kagan, P.M. Gallop, Specific detection of quinoproteins by redox-cycling staining., J. Biol. Chem. 266 (1991) 689–692. 10.1016/S0021-9258(17)35225-0.

[48] T. Bedada, Characterization of Tetrazolium Salts and Formazans using Computational Chemistry for Radiochromic Dosimetry, Electron. Thesis Diss. Repos. (2019). https://ir.lib.uwo.ca/etd/6458.

[49] M. Mompeán, V. Romano, D. Pantoja-Uceda, C. Stuani, F.E. Baralle, E. Buratti, D.V. Laurents, The TDP-43 N-terminal domain structure at high resolution, FEBS J. 283 (2016) 1242–1260. 10.1111/febs.13651.

[50] Q. Cao, D.R. Boyer, M.R. Sawaya, P. Ge, D.S. Eisenberg, Cryo-EM structures of four polymorphic TDP-43 amyloid cores, Nat. Struct. Mol. Biol. 26 (2019) 619–627. 10.1038/s41594-019-0248-4.

[51] Y. Porat, A. Abramowitz, E. Gazit, Inhibition of Amyloid Fibril Formation by Polyphenols: Structural Similarity and Aromatic Interactions as a Common Inhibition Mechanism, Chem. Biol. Drug Des. 67 (2006) 27–37. 10.1111/j.1747-0285.2005.00318.x.

[52] S.K. Sonawane, H. Chidambaram, D. Boral, N.V. Gorantla, A.A. Balmik, A. Dangi, S. Ramasamy, U.K. Marelli, S. Chinnathambi, EGCG impedes human Tau aggregation and interacts with Tau, Sci. Rep. 10 (2020) 12579. 10.1038/s41598-020-69429-6.

[53] H.J. Wobst, A. Sharma, M.I. Diamond, E.E. Wanker, J. Bieschke, The green tea polyphenol (−)-epigallocatechin gallate prevents the aggregation of tau protein into toxic oligomers at substoichiometric ratios, FEBS Lett. 589 (2015) 77–83. 10.1016/j.febslet.2014.11.026.

[54] S. Roy, R. Bhat, Suppression, disaggregation, and modulation of γ-Synuclein fibrillation pathway by green tea polyphenol EGCG, Protein Sci. 28 (2019) 382–402. 10.1002/pro.3549.

[55] J.D. Eaton, M.P. Williamson, Multi-site binding of epigallocatechin gallate to human serum albumin measured by NMR and isothermal titration calorimetry, Biosci. Rep. 37 (2017) BSR20170209. 10.1042/BSR20170209.

[56] T. He, Q. Liang, T. Luo, Y. Wang, G. Luo, Study on Interactions of Phenolic Acid-Like Drug Candidates with Bovine Serum Albumin by Capillary Electrophoresis and Fluorescence Spectroscopy, J. Solut. Chem. 39 (2010) 1653–1664. 10.1007/s10953-010-9608-8.

[57] A. Precupas, R. Sandu, A. Ruxandra Leonties, D.-F. Anghel, V. Tudor Popa, Complex interaction of caffeic acid with bovine serum albumin: calorimetric, spectroscopic and molecular docking evidence, New J. Chem. 41 (2017) 15003–15015. 10.1039/C7NJ03410E.

[58] D.D. Scott, D. Mowrey, K. Nagarajan, L. François-Moutal, A. Nair, M. Khanna, Molecular Dynamics simulation of TDP-43 RRM in the presence and absence of RNA, (2022) 2022.03.15.484514. 10.1101/2022.03.15.484514.

[59] J.C. Rengifo-Gonzalez, K. El Hage, M.-J. Clément, E. Steiner, V. Joshi, P. Craveur, D. Durand, D. Pastré, A. Bouhss, The cooperative binding of TDP-43 to GU-rich RNA repeats antagonizes TDP-43 aggregation, eLife. 10 (2021) e67605. 10.7554/eLife.67605.

