## Supplementary Figure-S1 for "*In silico* and *in vitro* studies suggest epigallocatechin gallate (EGCG), a polyphenol in green tea, can bind and modulate the aggregation and cytotoxicity of the full-length TDP-43 protein implicated in TDP-43 proteinopathies"

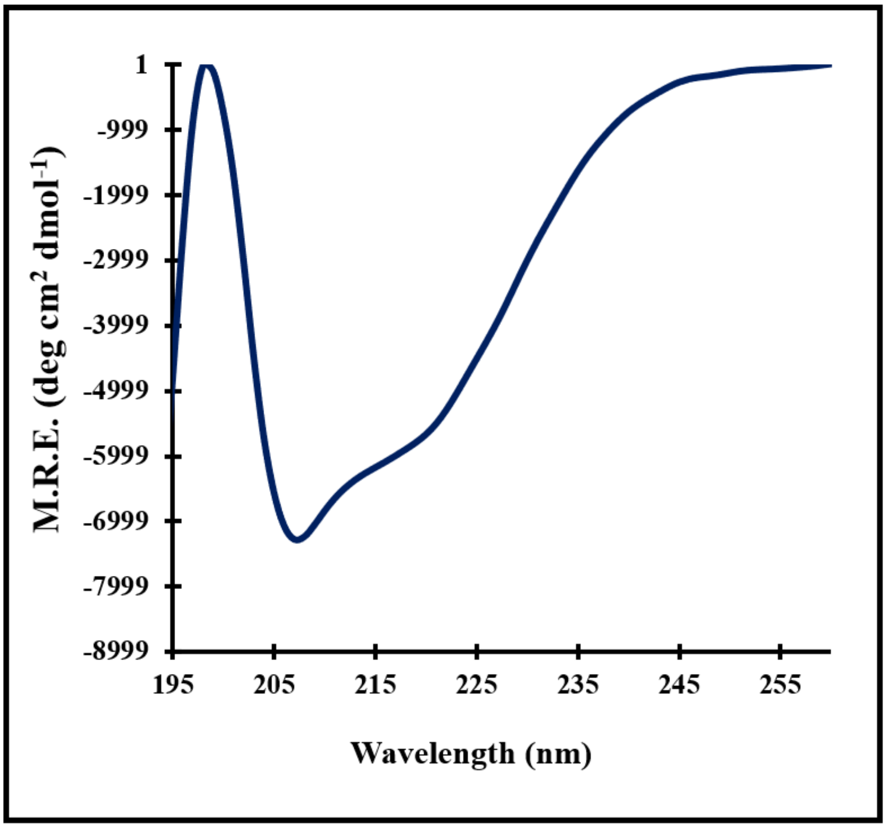


**Supplementary Figure S1: Far-UV Circular dichroism (CD) spectrum of refolded TDP-43.**

To analyse the secondary structure of TDP-43 in refolding buffer (0.3M Urea in 10mM Tris, pH 7) the far-UV CD spectrum was recorded using JASCO 1500 spectropolarimeter and 2mm path-length quartz cuvette at 25 $^{o}$C. The spectra were blank subtracted and converted to as Mean Residue Ellipticity, [ϴ]MRE, using the formula: [ϴ]MRE = (ϴ*MRW)/10*c*l, where ϴ represents ellipticity in millidegrees, c is concentration of protein in g/ml, l is path length in cm, and the mean residue weight (MRW) was taken as 112. The spectrum shows peaks and overall shape similar to as reported previously for the refolded TDP-43 thereby suggesting that our protein is also in a refolded conformation [1].

### Supplementary References


1. Vega, M.V., Nigro, A., Luti, S., Capitini, C., Fani, G., Gonnelli, L., Boscaro, F. and Chiti, F., 2019. Isolation and characterization of soluble human full-length TDP-43 associated with neurodegeneration. The FASEB journal, 33(10), pp.10780-10793.
