## Supplementary Figure-S2 for "*In silico* and *in vitro* studies suggest epigallocatechin gallate (EGCG), a polyphenol in green tea, can bind and modulate the aggregation and cytotoxicity of the full-length TDP-43 protein implicated in TDP-43 proteinopathies"

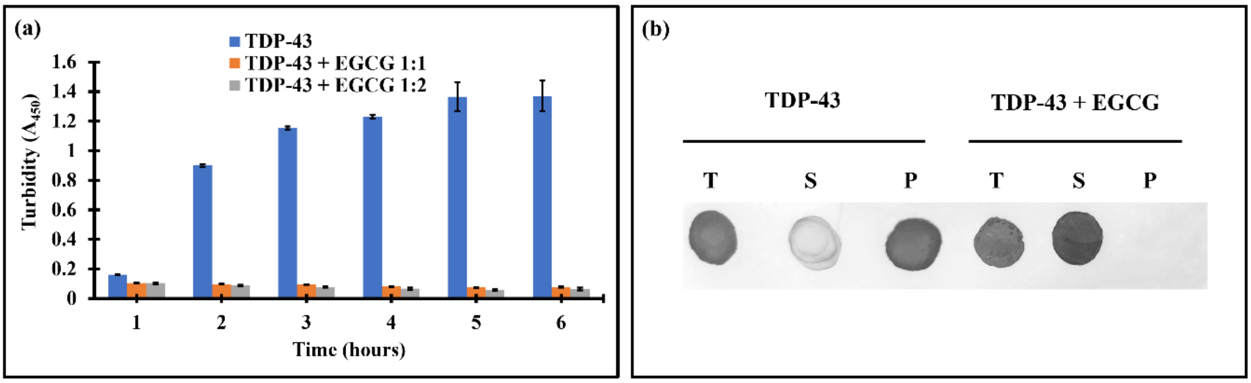


**Supplementary Figure-S2: EGCG inhibits the *in vitro* aggregation of TDP-43 in presence of 2.5M urea.**

**(a)** Changes in the turbidity of the TDP-43 solution (100μM) incubated in presence or absence of EGCG in aggregation buffer (500μL DTT, 2.5M urea in PBS, pH 7.5) was monitored by recording the absorbance at 450nm. The turbidity was recorded for all the TDP-43 samples incubated without or with EGCG at protein to EGCG stoichiometric ratios of 1:1 or 1:2 at 37$^{o}$C at an interval of every 1h upto 6h. **(b)** Sedimentation assay of TDP-43 samples aggregated in presence or absence of EGCG detected by spotting and staining on a PVDF membrane. TDP-43 aggregates prepared in presence or absence of 200μM EGCG in aggregation buffer (PBS pH 7.5, 2.5M urea and 500μM DTT) were centrifuged at 20,000× g for 45 minutes to separate supernatant with soluble protein from pellet containing the insoluble aggregated protein. Equal volumes of the unfractionated sample (total, T), the supernatant (S) and resuspended pellet (P) were added with 4× Laemmli reducing sample buffer, boiled and analysed by spotting and Coomassie staining on a PVDF membrane.
