## Supplementary Figure-S3 for "*In silico* and *in vitro* studies suggest epigallocatechin gallate (EGCG), a polyphenol in green tea, can bind and modulate the aggregation and cytotoxicity of the full-length TDP-43 protein implicated in TDP-43 proteinopathies"

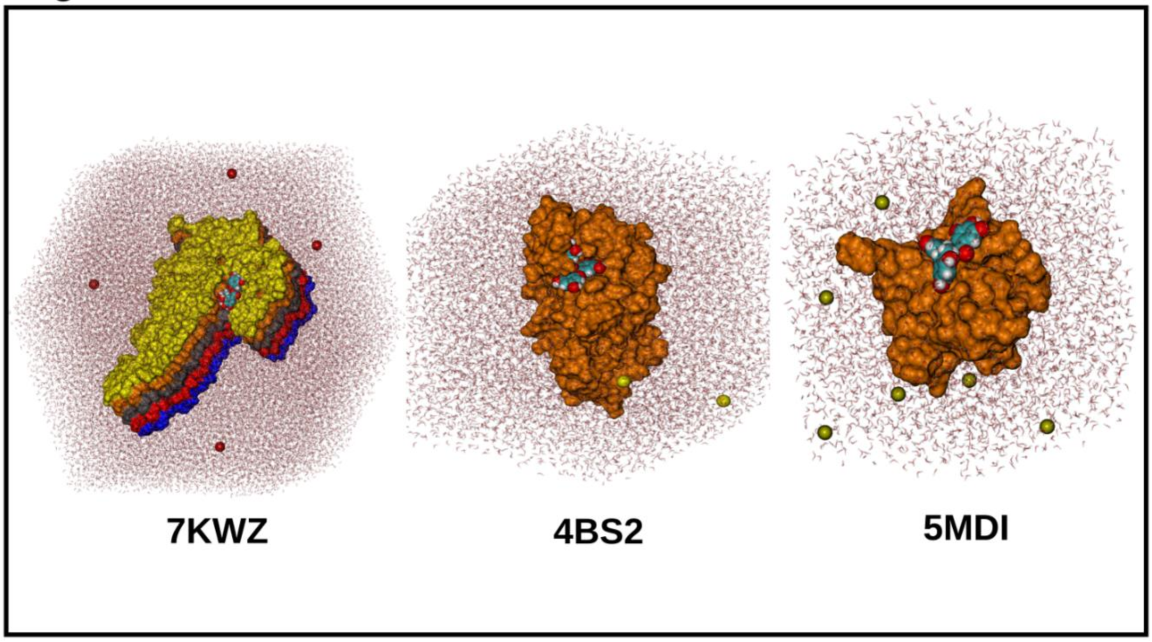


**Supplementary Figure S3: All-atom model of EGCG with the domain structures of TDP-43 with EGCG.**

**(a)** The ensemble cryo-EM structure of the low complexity C-terminal domain of TDP-43 (PDB ID: 7KWZ) (Chains A to E coloured in blue, red, silver, orange and yellow respectively) with EGCG (VDW representation). **(b)** The solution NMR structure of the RRM1-2 domain of TDP-43 (PDB ID: 4BS2) (Orange) with EGCG (VDW representation). **(c)** The crystal structure of N-terminal domain of TDP-43 (PDB ID: 5MDI) (Orange) with EGCG (VDW representation). Each structure is immersed in the box of water (represented as lines) and ions (red – $Cl^-$ ; yellow – $Na^+$ ).
