## Supplementary Figure-S4 for "*In silico* and *in vitro* studies suggest epigallocatechin gallate (EGCG), a polyphenol in green tea, can bind and modulate the aggregation and cytotoxicity of the full-length TDP-43 protein implicated in TDP-43 proteinopathies"

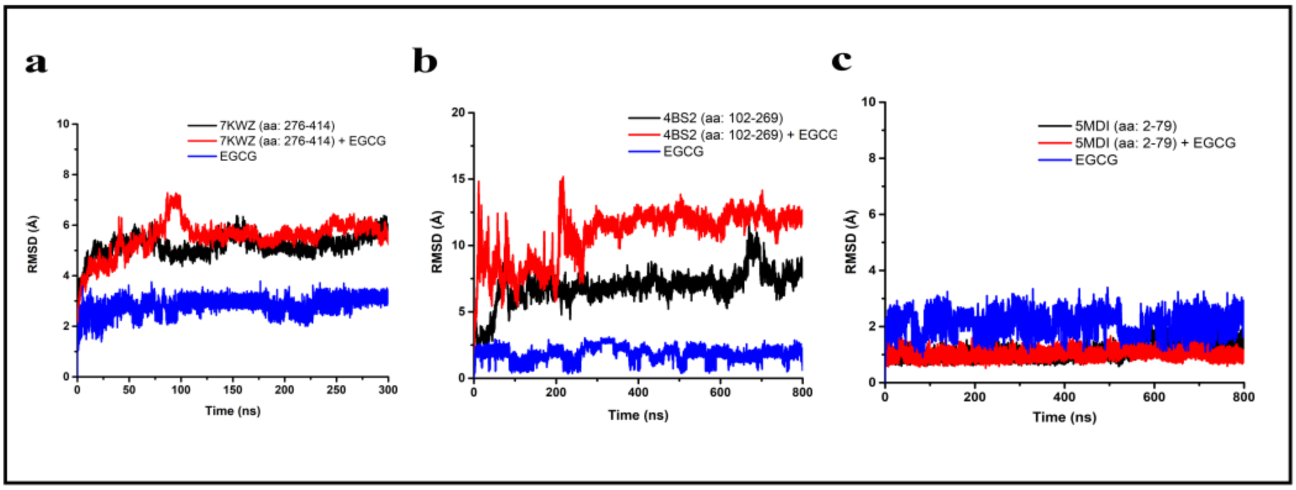


**Supplementary Figure S4: RMSD profile from the MD simulation of the domain structures of TDP-43 complexed with EGCG.**

**(a)** The C-α atom RMSD of the protein alone and protein EGCG simulation from the 300 ns simulation of the cryo-EM structure of the ensemble structure of the low complexity C-terminal domain of TDP-43 (PDB ID: 7KWZ, aa: 276-414) with EGCG along with the ligand alone RMSD. **(b)** The RMSD of the C-α atoms of the protein alone and protein-ligand simulation of 800 ns for the solution NMR structure of the RRM1-2 domain of TDP-43 (PDB ID: 4BS2, aa: 102-269) with EGCG along with the ligand alone RMSD from the simulation. **(c)** Root Mean Square Deviation (RMSD) of the protein backbone C-α atoms for the protein alone and the protein-ligand simulation of 800ns along with the ligand RMSD from the protein-ligand simulation for the N-terminal domain of TDP-43 (PDB ID: 5MDI, aa: 2-79).
