## Supplementary Table-S1 for "*In silico* and *in vitro* studies suggest epigallocatechin gallate (EGCG), a polyphenol in green tea, can bind and modulate the aggregation and cytotoxicity of the full-length TDP-43 protein implicated in TDP-43 proteinopathies"

**Supplementary Table S1: Components of the gmx_MMPBSA free energy calculation for the structures of the domains of TDP-43 with EGCG.**

| Energy Term | 5MDI (aa: 2-79) + EGCG (kcal/mol) | 4BS2 (aa: 102-269) + EGCG (kcal/mol) | 7KWZ (aa: 276-414) + EGCG (kcal/mol) |
|---|---|---|---|
| VDWAALS ($E_{VdW}$) | -3.92 ± 0.13 | -12.09 ± 0.08 | -37.21 ± 0.05 |
| EEL ($E_{ele}$) | -5.37 ± 0.23 | -10.02 ± 0.17 | -11.12 ± 0.08 |
| EPB ($E_{polar}$) | 7.78 ± 0.28 | 16.68 ± 0.18 | 31.51 ± 0.06 |
| ENPOLAR ($E_{non\ polar}$) | -0.54 ± 0.02 | -1.7 ± 0.01 | -3.47 |
| GGAS ($E_{VdW}$+$E_{ele}$) | -9.28 ± 0.33 | -22.92 ± 0.21 | -48.33 ± 0.08 |
| GSOLV ($E_{polar}$+$E_{non\ polar}$) | 7.24 ± 0.26 | 14.98 ± 0.18 | 28.04 |
| **Total ($E_{VdW}$+$E_{ele}$+$E_{polar}$+$E_{non\ polar}$)** | **-2.04 ± 0.09** | **-7.94 ± 0.07** | **-20.29 ± 0.06** |
